# Radial astroglia cooperate with microglia to clear neuronal cell bodies during zebrafish optic tectum development

**DOI:** 10.1101/2025.03.14.643334

**Authors:** Heather M. Barber, Cassidy G. Robbins, Zachary Cutler, Robin I. Brown, Inge Werkman, Sarah Kucenas

## Abstract

The clearance of dead cells by phagocytes is an essential component of neural development in many organisms. Microglia are the main phagocytes in the central nervous system (CNS), but the extent of participation by other glial cells remains unclear, especially under homeostatic conditions. During zebrafish optic tectum (OT) development, we observed radial astroglia forming dynamic, spherical projections from their basal processes. These projections, which we call scyllate heads, coincide with a wave of neuronal cell death in the OT. We show that scyllate heads surround the majority of dying neurons soon after phosphatidylserine exposure. However, unlike traditional phagosomes, scyllate heads persist for many hours and are rarely acidified or internalized. Instead, microglia invade scyllate heads and remove their contents for terminal degradation. Our study reveals an active role for radial astroglia in homeostatic cell clearance and cooperation between microglia and radial astroglia during zebrafish OT development.

**Highlights:**

- Optic tectum astroglia form large, dynamic projections called scyllate heads
- Scyllate heads surround the majority of dying neurons during a wave of apoptosis
- Scyllate heads are intermediate containers of dying cells rather than phagosomes
- Microglia invade scyllate heads to remove their contents for terminal degradation

## Introduction

During neural development, many cells undergo programmed cell death (PCD) and are efficiently cleared away by phagocytes^1,2^. The process by which phagocytes recognize, engulf, and degrade dead or abnormal cells is called efferocytosis^2,3,4,5^. Efferocytosis by professional phagocytes, including macrophages and microglia, has been extensively investigated^2,3,4,5^. However, studies demonstrate that many other cell types can participate in efferocytosis and are called non-professional phagocytes when acting in this context^3,4,6,7^. In phagocyte-deficient models, some non-professional phagocytes can increase their phagocytic activity to compensate for the lack of professional phagocytosis^6,7,8^. However, it is less understood if and how professional and non-professional phagocytes cooperate to clear debris during physiological development.

One of the most well-known examples of developmental PCD occurs during neurogenesis, when neurons unnecessary for circuit function undergo PCD and are cleared by microglia, the resident phagocytes of the central nervous system (CNS)^2,5,9^. In the zebrafish optic tectum (OT), the major visual processing center in zebrafish, retinal ganglion cell axons that form the optic nerve begin to connect to OT targets at approximately 60 hours post fertilization (hpf)^10,11^. At this same stage, there is a sharp increase in the number of dead and dying OT neurons^2,12^. Microglia, which originate outside of the CNS, begin infiltration into the OT around 60 hpf and do not reach their stable larval population size until 72 to 96 hpf^12,13,14^. Thus, there are few professional phagocytes compared to the large number of dying cells and debris present during this critical developmental window. Previously, we showed that during early zebrafish development when microglia and macrophages have not yet colonized the zebrafish trunk, neural crest cells are able to phagocytose debris^15^. Therefore, we hypothesized that another non-professional phagocytic cell population contributes to efferocytosis in the developing OT.

In the OT, there is a population of glial cells called radial astroglia which exhibit astrocyte-like properties such as blood-brain barrier control and association with synapses^16,17,18^. Astrocytes and astrocyte-like cells can function as non-professional phagocytes^2,8,19,20^ and in mice, astrocytes assist microglia in the clearance of neurite debris as well as respond to dying cells when microglial activity is impaired^21^. Therefore, we hypothesized that OT radial astroglia participate in neuronal clearance during this developmental window of high phagocytic load and low microglial activity.

In this study, we utilized live imaging to investigate the role of OT radial astroglia in the clearance of dying cells. We demonstrate that during physiological zebrafish development, OT radial astroglia form large spherical structures from their basal processes that surround and hold cell bodies, particularly those of dying neurons. These structures, which we call scyllate heads, are highly dynamic, often long-lasting, and rarely become acidic, distinguishing them from classically described phagosomes. Microglia subsequently interact with scyllate heads to remove contents from these compartments and complete the degradation process. Together, our findings uncover a previously undescribed role for radial astroglia in the OT and reveal highly coordinated interactions between professional and non-professional phagocytes.

## Results

### OT radial astroglia form scyllate heads during a critical developmental stage

In the zebrafish OT, the axons of retinal ganglion cells (RGCs) begin to innervate at approximately 60 hours post fertilization (hpf)^10,11^. Concurrently, there is an increase in OT neuronal death between 60 and 72 hpf^2,12,14^. To observe how radial astroglia behave during this critical point in development, we imaged the OT between 55 and 84 hpf for 6 to 12 hours in *Tg(slc1a3b:myrGFP-P2A-H2AmCherry)* larvae from a sagittal view (Figure 1A schematic, Figure S1A). This transgene uses regulatory sequences for *slc1a3b,* which encodes the protein glutamate astrocyte specific transporter (GLAST), an established marker for astroglia, to drive expression of GFP in astroglial membranes and mCherry in astroglial nuclei^22^. In these time-lapses, we observed OT radial astroglia form large spherical structures, which emerged from small branches off their basal processes (Figure 1A; Video S1). In our imaging studies, we observed many of these membranous structures appearing *en masse*, often moving around dynamically at the end of thin, neck-like projections before dissipating. Due to their striking morphology and the consistency at which we observed them in nearly all our time-lapses, we sought to investigate the nature of these radial astroglial structures.

**Figure 1.**
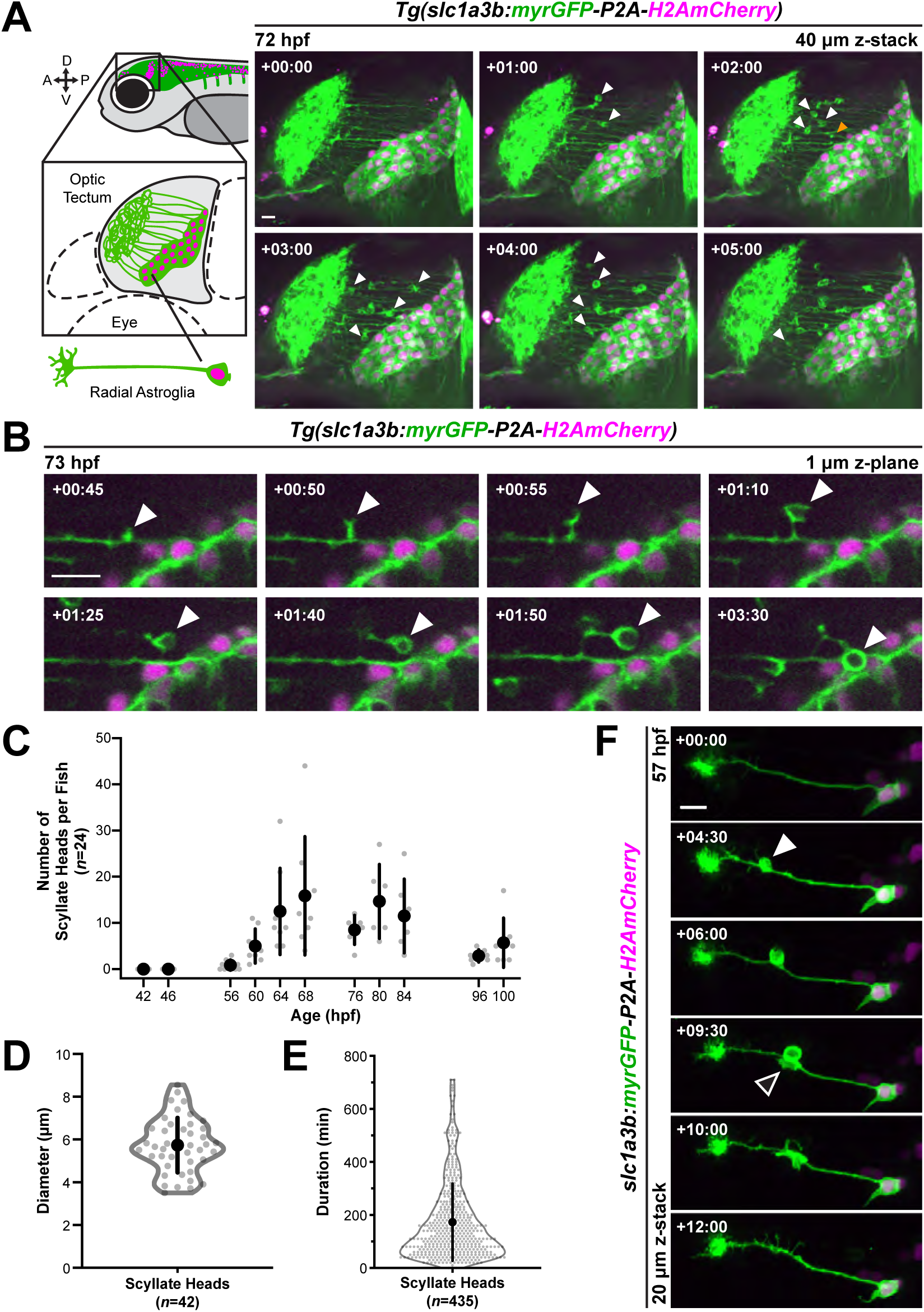
OT astroglia form scyllate heads from their basal projections during a critical developmental period. A) Schematic depicts area and orientation of imaging, pattern of transgenic labeling, and example of a single radial astroglia. 40-µm projection stills taken from a time-lapse from 72 to 77 hpf of a *Tg(slc1a3b:myrGFP-P2A-H2AmCherry)* larva where astroglial membranes (green) and nuclei (magenta) are labeled. Arrowheads indicate recently formed scyllate heads. B) Enlarged single z-plane images of an individual scyllate head from A (orange arrowhead). C) Quantification of the number of scyllate heads throughout early development from time-lapses of *Tg(slc1a3b:myrGFP-P2A-H2AmCherry)* larvae, *n*=24 fish. D) Quantification of scyllate head diameter. Mean diameter=5.73±1.3 µm, *n*=42 scyllate heads from 3 fish. E) Quantification of scyllate head duration. Average duration=173±146 min, median duration=130 min, range: 0 min (1 time-point) to 710 min (entire time-lapse), *n*=435 scyllate heads from 9 fish. F) 20 μm projection stills taken from a time-lapse of a single radial astroglia obtained from sparse labeling with *slc1a3b:myrGFP-P2A-H2A-mCherry*, beginning at 57 hpf. The astroglia (green membranes, magenta nucleus) forms a scyllate head (arrowhead) which later dissipates (arrowhead outline). Time indicated in hh:mm after start of imaging. Scale bars, 10 μm.

To be sure that the emergence of these structures was not a result of the transgene itself, we confirmed that they were present in other transgenic lines. We generated three stable *slc1a3b*-driven transgenic lines: a myristoylated membrane-bound *Tg(slc1a3b:EGFP)*, a CAAX-localized membrane-bound *Tg(slc1a3b:TagRFP-CAAX)*, and a cytoplasmic *Tg(slc1a3b:mCerulean3)*. We then imaged larvae of each transgenic line in time-lapses between 55 and 84 hpf (Figure S1B). In all these transgenic lines, we observed the same structures formed by radial astroglia and sought to investigate their function.

Early in development, radial glia act as neural and glial progenitors^17,18^, so we considered the possibility that these structures contained the recent progeny of radial astroglia. The *Tg(slc1a3b:myrGFP-P2A-H2AmCherry)* transgene labels both membranes (myrGFP) and nuclei (H2AmCherry) of astroglial cells^22^. The spherical structures observed in our time-lapses did not contain the mCherry nuclear marker, indicating that they were not astroglial cell bodies (Figure S1C). Additionally, these structures usually emerged from the basal process near the middle of the OT rather than the area near the ventricle where the astroglial cell bodies reside. Most notably, we did not observe astroglial nuclei dividing during our time-lapses. Therefore, we conclude that these structures were not a result of astroglial proliferation.

Due to the known role of astroglia in blood-brain barrier control^16,17,18^, we also investigated the possibility that these structures were interacting with blood vessels. We imaged *Tg(slc1a3b:TagRFP-CAAX);Tg(fli1a:EGFP)* transgenic larvae from 55 hpf to 67 hpf to visualize both astroglial membranes and blood vessels. We determined that, while astroglial membranes did surround blood vessels in tube-like formations, the spherical structures we observed were not associated with blood vessels (Figure S1D).

After these initial investigations and thorough examination of existing literature, we conclude that our observations represent a previously undescribed behavior of OT radial astroglia. To discuss these structures while also avoiding incorrectly labeling them as existing structures, we decided to give them a new name. Due to the way that many of the structures suddenly form as mouth-like cups on long neck-like branches, we decided to call them “scyllate heads” after Scylla, a cliff-dwelling, many-headed monster in Homer’s Odyssey who eats sailors from passing ships. For the purposes of our study, we defined scyllate heads as spherical or hemi-spherical (cup-like) structures formed by radial astroglial membranes.

### Scyllate head formation and dissipation is dynamic and highly variable

To begin to characterize scyllate heads, we analyzed *Tg(slc1a3b:myrGFP-P2A-H2AmCherry)* larvae in 6 or 12 hour time-lapses between 54 and 86 hpf. For most of our imaging, we focused on the most lateral 40 µm of the OT, the lateral edge of which we identified as the first z-plane in which we clearly identified astroglial basal processes (Figure S1A). However, we did observe scyllate heads deeper in the OT (Figure S1E), indicating that this phenomenon is not limited to a specific OT layer or region.

The number of scyllate heads present varied by the age of the larvae. To create a timeline of scyllate head formation, we counted the number of scyllate heads over time between 42 and 100 hpf, beginning after OT expression of the *Tg(slc1a3b:myrGFP-P2A-H2AmCherry)* transgene, which starts at approximately 40 hpf (Figure 1C). From this analysis we found that the peak of scyllate head formation was approximately 68 hpf and gradually decreased over the next 24 hours (Figure 1C). By 100 hpf, we rarely observed scyllate heads.

In our imaging, we observed a high amount of variation in scyllate head characteristics. We found that the average diameter of a scyllate head was 5.73±1.27 µm (Figure 1D) which is similar to the average size of a nucleus^23^. We also assayed how long scyllate heads were maintained after formation by determining the time-point of a scyllate head’s first appearance and of its dissipation, the difference in which we call the duration. From our characterization, the average duration of a scyllate head was 172±146 minutes (Figure 1E). The duration ranged from 0 minutes, or 1 time-point, to 710 minutes, the entire time-lapse, with a median duration of 130 minutes (Figure 1E). Occasionally, we observed scyllate heads that were already present when we began imaging or had not yet dissipated by the end of imaging, meaning that the measured duration is shorter than the actual duration for these scyllate heads. We conclude that scyllate heads vary in duration, with a noticeable skew towards lower durations, but can be long-lasting.

To better visualize scyllate head dynamics, we injected the *slc1a3b:myrGFP-P2A-H2AmCherry* plasmid into embryos to mosaically label astroglia. We then captured the emergence of scyllate heads from the basal processes of single astroglia (Figure 1F, Video S2). In this representative time-lapse beginning at 57 hpf, we observed small projections emitting from a main astroglial basal process followed by the formation of a scyllate head at 62 hpf (+4:30). The scyllate head was present for 320 minutes before suddenly dissipating at 67 hrs (+9:50). The method of dissipation of this scyllate head was surprising, as it appeared to burst open before reincorporating into the basal process, rather than being transported to the cell body or pruned by another cell. We saw this method of dissipation in other examples of scyllate heads, including in stable lines. We also observed some dissipating scyllate heads which appeared to open and collapse back into the membrane more gradually, and still others which suddenly enlarged significantly before similarly collapsing.

Mosaic labeling of astroglia also allowed us to make direct observations about the behavior of individual scyllate heads. We saw that some scyllate heads appeared to be directly in contact with the basal process (Figure 1F) while others formed on the ends of thin neck-like branches that protruded from the basal process (Figure 1B, Video S2, Figure S1F). We often imaged radial glia that did not form any scyllate heads during the imaging window (data not shown). In contrast, we imaged astroglia that formed and maintained multiple scyllate heads simultaneously during the imaging window (Figure S1F). These findings indicate that scyllate head formation is not a one-to-one process and is highly dynamic and variable. We therefore hypothesized that scyllate heads are formed in response to an extrinsic cue.

### RGC axon infiltration and visual input are dispensable for scyllate head formation

Scyllate head formation occurs in the OT during the window in which RGCs are creating synaptic connections with OT neurons. Therefore, we considered the possibility that vision and/or RGC axon infiltration may play a role in scyllate head formation. To determine if the presence of RGC axons (and subsequent visual input) affected scyllate head formation, we utilized *lakritz (lak)* mutant fish. *Lak* mutants have a missense mutation in *atonal bHLH transcription factor 7 (atoh7,* also called *ath5)*, a transcription factor important for RGC differentiation^24,25^. Without *atoh7* function, RGC axons do not develop and do not infiltrate the OT and therefore, *lak* mutant larvae are completely blind^24^. We imaged *Tg(atoh7:mRFP);Tg(slc1a3b:myrGFP-P2A-H2AmCherry)* larvae from 51 hpf to 67 hpf to capture the stage when RGCs infiltrate the OT and scyllate heads begin to form. In *lak* heterozygous larvae, we observed RGC axons innervating the OT (Figure 2A, Video S3). In *lak* homozygous mutant larvae, while we did see mRFP expression in the retina, we saw no RGC axons (Figure 2A, Video S3). Despite the lack of RGC axon infiltration and vision, we still observed scyllate head formation in *lak* homozygous mutant larvae (Figure 2A, Video S3) and there was no apparent difference in the appearance, timing, or number of scyllate heads between heterozygous and homozygous mutant *lak* sibling larvae (Figure 2A’). Therefore, though RGC axon infiltration and scyllate head formation occur in the same developmental window, we conclude that RGC axon infiltration and visual input are dispensable for scyllate head formation.

**Figure 2.**
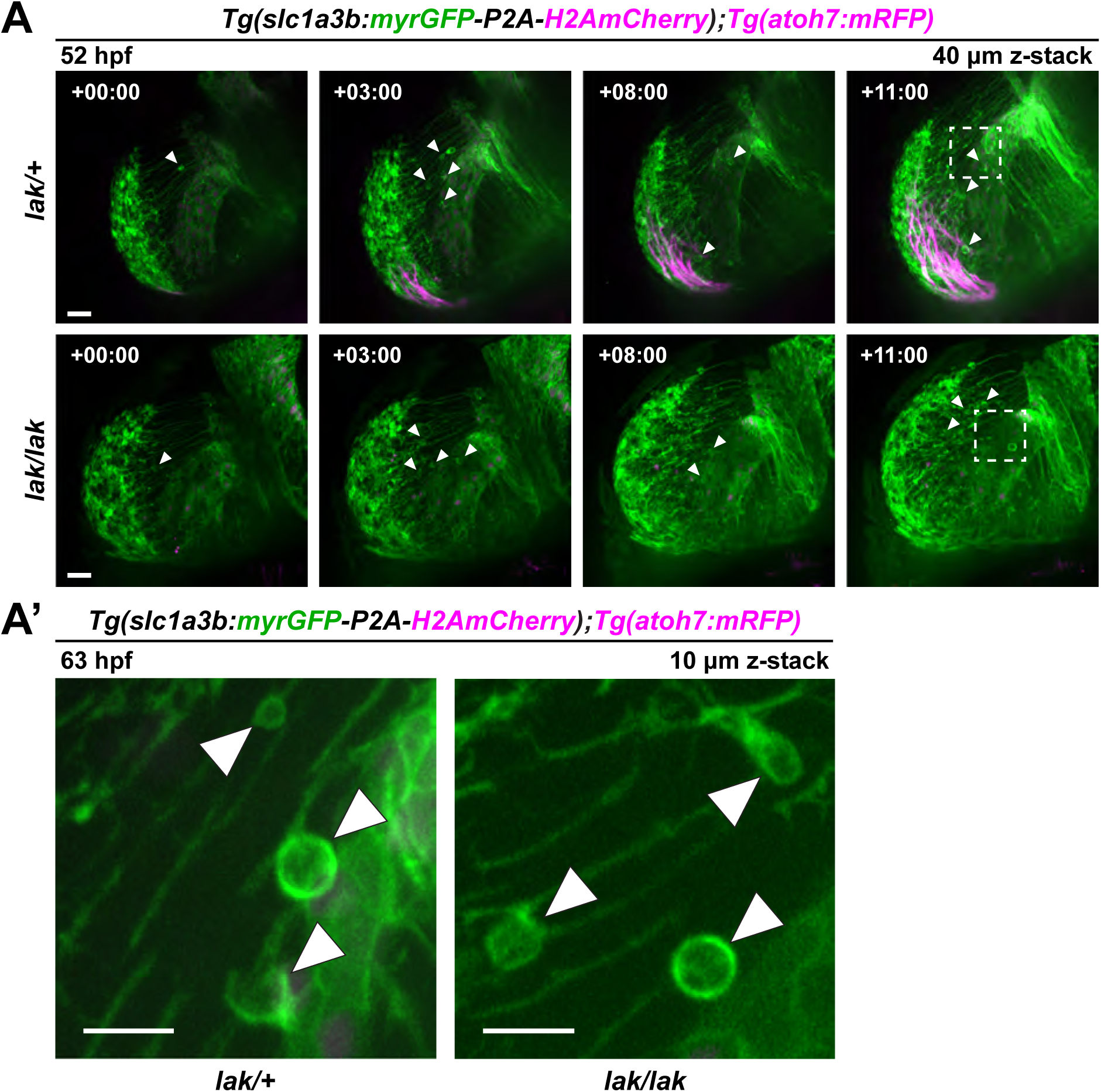
Input of retinal ganglion cell axons into the optic tectum is dispensable for scyllate head formation. A) Still images from time-lapses of 52 hpf *Tg(slc1a3b:myrGFP-P2A-H2AmCherry);Tg(ath5:mRFP)*;*lakritz (lak)* heterozygous (top) and homozygous (bottom) larvae. Images shown are 40 µm projections. Astroglia (green membranes, magenta nuclei) and RGC axons (magenta) are labeled. White arrowheads indicate recently formed scyllate heads. A’) Insets showing 10 µm projections taken from A at 60 hpf, indicated by dashed boxes. Time indicated in hh:mm after start of imaging. Scale bars, 20 µm (A), 10 µm (A’).

### Scyllate heads contain the cell bodies of dying neurons

In our imaging, we observed that scyllate heads often began as cup-like structures that eventually became fully enclosed, much like phagocytic cups made by professional phagocytes. Therefore, we hypothesized that scyllate heads surrounded and perhaps even engulfed something in the environment. Because scyllate heads are generally large enough to accommodate a cell body (Figure 1D), we looked at other cells as the potential contents. Major cell types in the OT between 60 and 72 hpf include oligodendrocyte progenitor cells, neurons, and microglia^26,27^. There is a large increase in PCD of neurons at 60 hpf^12,14^, the same stage that scyllate heads begin to form, and previous studies show that astroglia interact with neurons in various ways^16,28,29^. Therefore, we hypothesized that scyllate heads contained neurons. To investigate this possibility, we generated the transgenic line *Tg(elavl3:H2B-GCaMP6s)*, in which *elavl3^+^* neurons express the calcium indicator GCaMP6s localized to the nucleus, where fluorescence intensity increases when calcium concentration increases^30^. Alongside this neuronal transgene, we labeled astroglial membranes with *Tg(slc1a3b:TagRFP-CAAX)* and imaged from 60 to 84 hpf in 6 or 12 hour time-lapses. Intriguingly, we observed that scyllate heads often contained bright GCaMP^+^ neuronal nuclei (Figure 3A, S2A). We also observed scyllate heads surrounding neurons with dim GCaMP signal that eventually became bright (GCaMP^+^) while within the scyllate head during our time-lapse imaging (Figure S2B). Of the scyllate heads that formed in our movies, 87.0±8.4% contained GCaMP^+^ neuronal nuclei (*n*=6 fish, Figure 3B). Therefore, we conclude that scyllate heads surround and contain OT neuronal soma.

**Figure 3.**
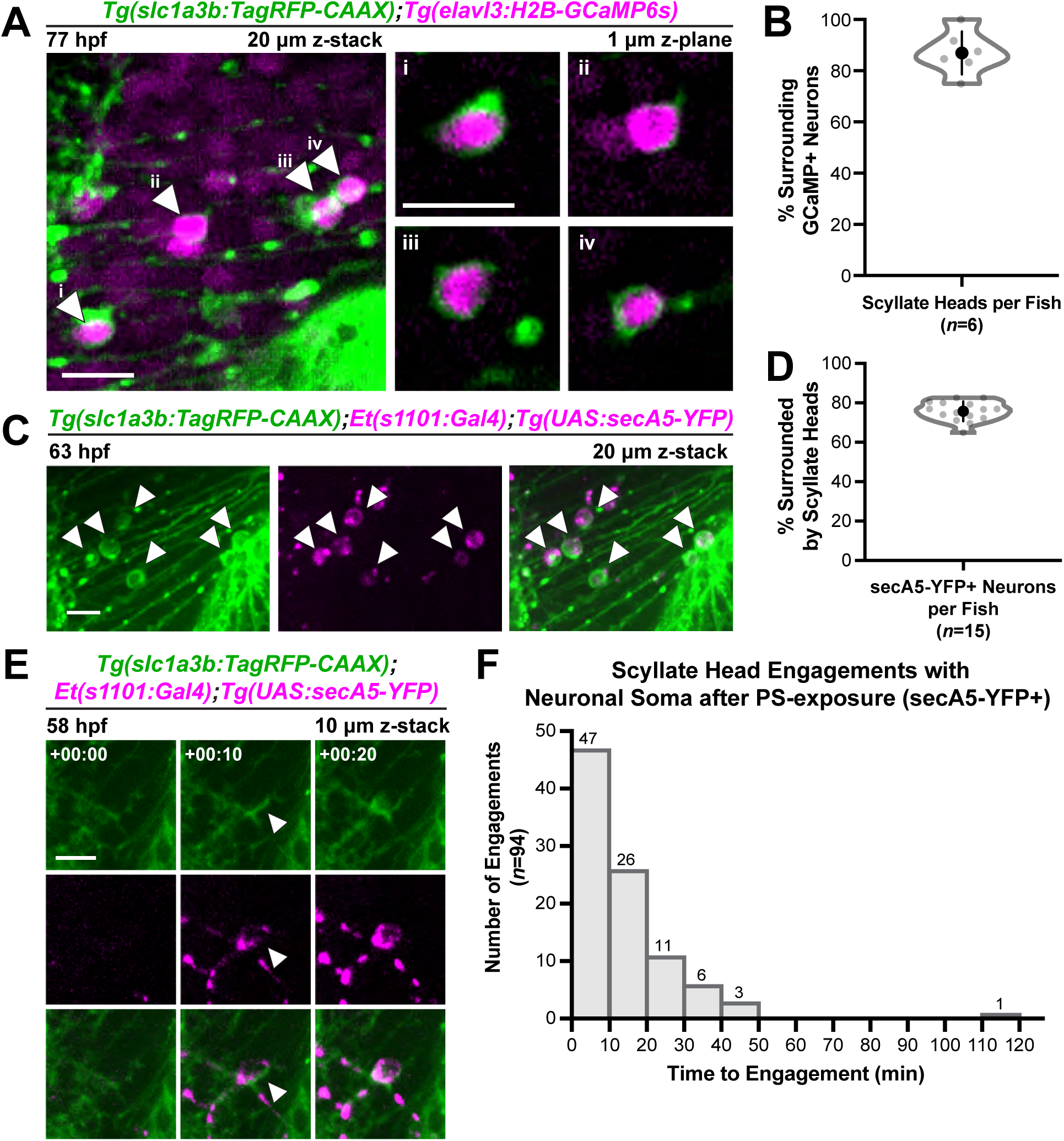
Scyllate heads surround dying neurons. A) 20 µm projection image of scyllate heads (green) surrounding the nuclei of *elavl3*^+^ neurons (magenta) in a 77 hpf *Tg(slc1a3b:TagRFP-CAAX);Tg(elavl3:H2B-GCaMP6s)* larva. Scyllate heads are indicated with white arrowheads. Insets show 1 µm z-planes of corresponding scyllate heads. Inset scale bar, 5 µm. B) Quantification of the percent of scyllate heads with GCaMP^+^ contents from time-lapses of *Tg(slc1a3b:TagRFP-CAAX)*;*Tg(elavl3:H2B-GCaMP6s)* larvae from 60 hpf to 84 hpf. Mean=87.0±8.4%, *n*=6 fish. C) Scyllate heads (green) surrounding dying neuronal soma (magenta) in a 63 hpf *Tg(slc1a3b:TagRFP-CAAX);Et(s1101:Gal4);Tg(UAS:secA5-YFP)* larva, 20 µm projection. Dying neurons surrounded by scyllate heads are indicated with white arrowheads. D) Quantification of the percent of dying OT neuronal soma (secA5-YFP^+^) surrounded by scyllate heads during time-lapses of *Tg(slc1a3b:TagRFP-CAAX);Et(s1101:Gal4);Tg(UAS:secA5-YFP)* larvae from 55 hpf to 83 hpf. Mean=75.7±5.0%, *n*=15 fish. E) 10 µm projection stills from a time-lapse of a *Tg(slc1a3b:TagRFP-CAAX);Et(s1101:Gal4);Tg(UAS:secA5-YFP)* larva showing a scyllate head (green, arrowhead) engagement with a secA5-YFP^+^ neuron (magenta). Images were taken every 10 minutes starting at 54 hpf. Time indicated in hh:mm after start of imaging. F) Quantification of the time from when *s1101^+^* neuronal PS exposure (secA5-YFP^+^) to scyllate head engagement during time-lapses of *Tg(slc1a3b:TagRFP-CAAX);Et(s1101:Gal4);Tg(UAS:secA5-YFP)* larvae from 54 hpf to 83 hpf. *n*=94 neurons from 9 fish. Scale bars, 10 µm.

During our time-lapses of *Tg(slc1a3b:TagRFP-CAAX);Tg(elavl3:H2B-GCaMP6s)* larvae from 60 to 84 hpf, we also observed scyllate heads surround neuronal nuclei which shrunk during the imaging window (Figure S2C). We suspected that this apparent nuclear shrinkage could be pyknosis, also known as chromatin condensation, a phenomenon where a nucleus condenses during cell death^31,32,33^. Additionally, changes in calcium level, potentially like what we observed when scyllate head contents suddenly became GCaMP^+^ (Figure S2B), are also associated with cell death^33,34^. During OT development, there is an increase in neuronal PCD^2,12^ concurrent with the period of scyllate head formation (Figure 1C). Therefore, we hypothesized that the neurons held by scyllate heads were dead or dying. To confirm that the nuclear shrinkage we observed was indeed pyknosis by apoptotic cells, we live-stained *Tg(slc1a3b:TagRFP-CAAX)* larvae at 72 hpf with the DNA-labeling dye Acridine Orange (AO) and imaged immediately following treatment (Figure S2D). AO is used to label condensed DNA found in early-stage apoptotic cells, indicated by bright AO fluorescence^35^. In these larvae, we observed scyllate heads with contents that were AO^+^ (Figure S2D). Of the scyllate heads present at the time of imaging, 41.9±16.9% held AO^+^ contents (Figure S2E). This is consistent with our previous observations of GCaMP^+^ neurons undergoing nuclear shrinkage after being surrounded by scyllate heads (Figure S2B) and led us to hypothesize that scyllate heads may surround cells in response to an early apoptotic cue. Overall, this data shows that scyllate heads can contain apoptotic cells but does not confirm that all scyllate head contents are dying. We thus further investigated the type of contents held by scyllate heads.

To directly show that scyllate heads surround dying neurons, we imaged *Tg(slc1a3b:TagRFP-CAAX);Et(s1101:Gal4);Tg(UAS:secA5-YFP)* larvae from 60 to 84 hpf in 6 or 12 hr time-lapses. The *s1101* enhancer trap line labels many neurons, including a large subset of OT neurons at 60 hpf^35,36^. This line drives expression of a secreted annexin V (secA5), which binds to phosphatidylserine (PS) exposed on the cell body of dying cells^37^, thus effectively labeling dead or dying *s1101^+^* neurons (secA5-YFP^+^). In our imaging, we observed scyllate heads surrounding secA5-YFP^+^ neuronal cell bodies (Figure 3C, Video S4). Of the neuronal soma that became secA5-YFP^+^, 75.7±5.0% were surrounded by scyllate heads during our 12-hour time-lapses (Figure 3D). This data demonstrates that astroglia interact with the majority of dying neurons by surrounding them in scyllate heads before they are cleared.

To determine how quickly radial astroglia recognized dying neurons, we quantified the time from when a neuron became secA5-YFP^+^ to when it was engaged by a scyllate head, which we defined as when a cup-like projection from a radial astroglia contacted a neuron that it would eventually surround during the time-lapse. From our imaging, we observed that in most events, neurons were engaged by a scyllate head less than 10 minutes after becoming secA5-YFP^+^ (47/94), and in almost all cases (93/94) were engaged within 1 hour (*n*=94 engagement events from 8 fish, Figure 3E-F). Together, these data demonstrate that scyllate heads respond robustly and quickly to neuronal cell death.

### Scyllate heads rarely acidify or bring contents to radial astroglial cell bodies for degradation

Our data demonstrates that scyllate heads respond to and surround dying neurons. Therefore, we sought to investigate whether they were directly involved in phagocytic clearance of these neurons. In several studies, engulfment by a phagocyte is defined as debris being internalized into the cell body^7,39^. Therefore, in time-lapses of *Tg(slc1a3b:TagRFP-CAAX)Et(s1101:Gal4);Tg(UAS:secA5-YFP)* larvae from 60 to 84 hpf, we tracked secA5-YFP^+^ neuronal cell bodies that were surrounded by scyllate heads. We defined the scyllate head contents as “internalized” when we observed the secA5-YFP^+^ debris in contact with an astroglial cell body without a boundary formed by the scyllate head (see Figure 4A). We found that only 4.8±5.9% of neuronal debris within scyllate heads was transported to the radial astroglial cell body and internalized (*n*=15 fish, Figure 4B). Most scyllate heads remained distant from the cell body.

**Figure 4.**
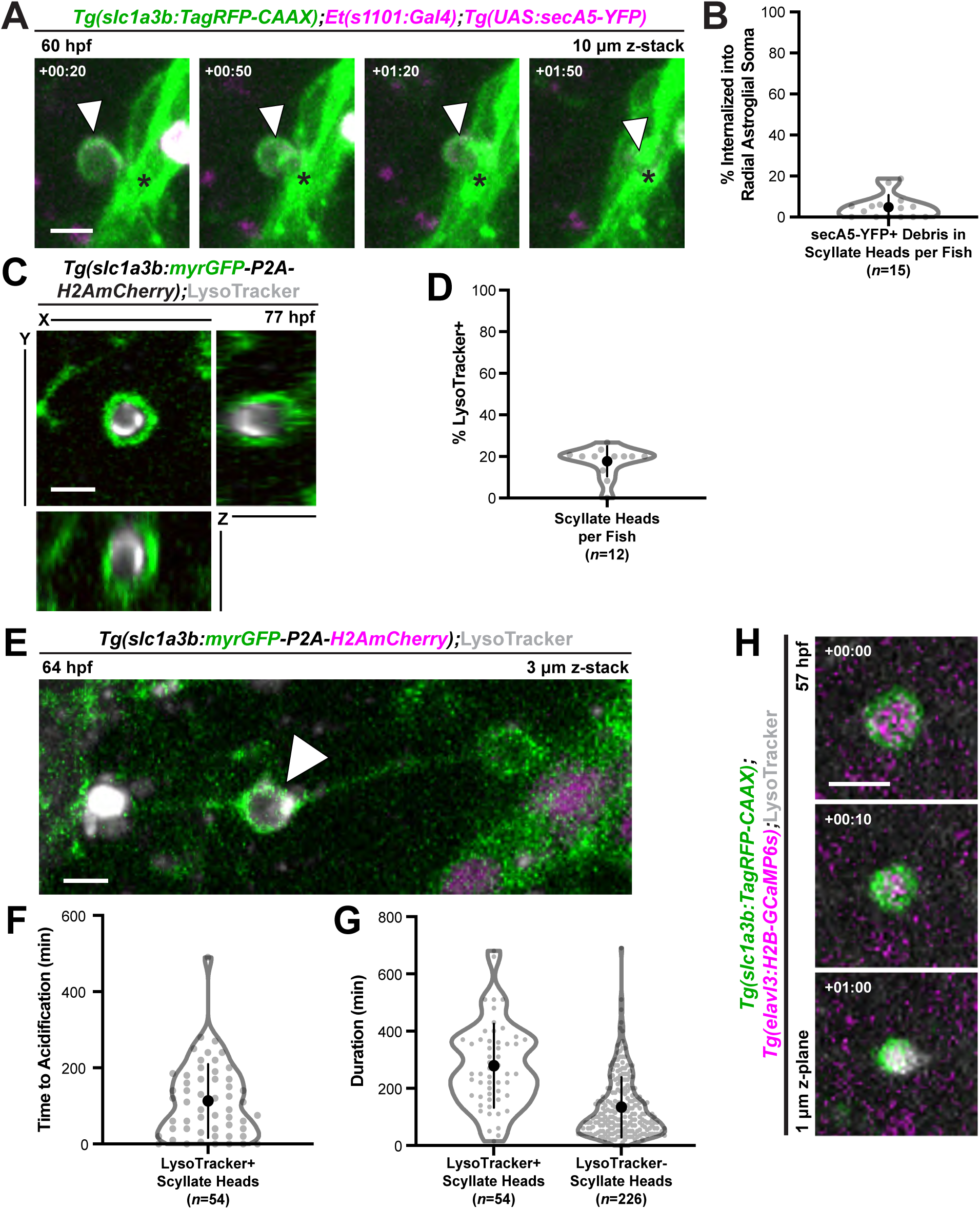
Scyllate heads can acidify while holding neuronal cell bodies but do so rarely. A) 10 µm projection stills from a time-lapse of a *Tg(slc1a3b:TagRFP-CAAX);Et(s1101:Gal4);Tg(UAS:secA5-YFP)* larva at 60 hpf. SecA5-YFP^+^ neuronal debris (magenta) contained in a scyllate head (green, arrowhead), is internalized into an astroglial soma (green, asterisk). B) Quantification of the percent of secA5-YFP^+^ neuronal debris in scyllate heads that were internalized by astroglia during time-lapses of *Tg(slc1a3b:TagRFP-CAAX);Et(s1101:Gal4);Tg(UAS:secA5-YFP)* larvae between 54 and 83 hpf. Average=4.8±5.9%, *n*=15 larvae. C) Isometric view of a scyllate head (green) from a 77 hpf *Tg(slc1a3b:myrGFP-P2A-H2AmCherry)* larva treated with 10 µM LysoTracker (white). D) Quantification of the percent of scyllate heads that become LysoTracker^+^ during time-lapses of *Tg(slc1a3b:TagRFP-CAAX);Tg(elavl3:H2B-GCaMP6s)*or *Tg(slc1a3b:myrGFP-P2A-H2AmCherry);Tg(mfap4:mTurquoise)* LysoTracker-treated larvae between 54 and 85 hpf. Average=17.7±7.2%, *n*=12 fish. E) 3 µm projection image of a scyllate head (green, arrowhead) positive for LysoTracker (white) in a LysoTracker-treated *Tg(slc1a3b:myrGFP-P2A-H2AmCherry)* larva at 64 hpf. The scyllate head remains along the radial astroglia’s basal process (green) away from the cell bodies (magenta). F) Quantification of the time to acidification of LysoTracker^+^ scyllate heads, defined at the difference between scyllate head formation and the time of first visible LysoTracker^+^ contents, from LysoTracker-treated larvae, see D. Average=113±97 minutes, n=54 scyllate heads from 12 fish. G) Quantification of scyllate head duration separated by LysoTracker^+^ and LysoTracker^-^ scyllate heads from LysoTracker-treated larvae, see D. Averages=279±146 minutes vs 134±105 minutes, *n*=54 LysoTracker^+^ and 226 LysoTracker^-^ scyllate heads from 12 fish. H) Single z-plane stills from a time-lapse of a *Tg(slc1a3b:TagRFP-CAAX);Tg(elavl3:H2B-GCaMP6s)* larva treated with LysoTracker beginning at 57 hpf. The scyllate head (green) holds a GCaMP^+^ neuron (magenta) before becoming LysoTracker^+^ (white). Time indicated in hh:mm after start of imaging. Scale bars, 5 µm.

Another important aspect of efferocytosis is the degradation of contents via phagolysosomal pathways^4,32^. A hallmark of degradation is the acidification of phagolysosomes, so we investigated whether scyllate heads could acidify. To do this, we used LysoTracker, a pH indicator often used to label lysosomes and phagolysosomes^40^. We treated either *Tg(slc1a3b:TagRFP-CAAX);Tg(elavl3:H2B-GCaMP6s)*or *Tg(slc1a3b:myrGFP-P2A-H2AmCherry);Tg(mfap4:mTurquoise)* larvae at 54 or 72 hpf with 10 μM LysoTracker Deep Red for 1 hour, then imaged for 6 or 12 hours immediately after treatment. In these studies, we observed that only a small subset of scyllate heads became Lysotracker^+^ (17.7±7.2%, *n*=12 fish, Figure 4C-D). Of these acidified scyllate heads, only a few (9/54 scyllate heads from 12 fish) moved to the astroglial soma for internalization. Most acidified scyllate heads became and remained acidic while distant from the cell body, as shown in Figure 4E.

The majority of the scyllate heads we tracked remained negative for LysoTracker in our imaging studies. On average, those that did become LysoTracker^+^ did so within 113±97 minutes of observed formation, with the time to acidification ranging from 0 minutes to 490 minutes (*n*=54 scyllate heads from 12 fish, Figure 4F). Additionally, we observed non-acidified scyllate heads that were long-lasting, with durations up to 690 minutes (Figure 4G). This demonstrates that acidification does not necessarily depend on how long the scyllate head contains its cargo. On average, the total duration of LysoTracker^+^ scyllate heads (*n*=54) was 279±146 minutes compared to 134±105 minutes in LysoTracker^-^ scyllate heads (*n*=226) in the same fish (Figure 4G). Additionally, a greater proportion of acidified scyllate heads were present at the end of the time-lapse compared to non-acidified scyllate heads (48% vs 18%, *n*=54 and 266 scyllate heads from 12 fish, respectively). Overall, acidified scyllate heads lasted longer and dissipated less often in our time-lapses, suggesting that scyllate heads are not effective at degrading contents and do not usually dissipate via this method.

We also considered the possibility that LysoTracker^+^ scyllate heads were surrounding non-cell body debris that they may be better equipped to degrade. We further analyzed our time-lapses of LysoTracker-treated *Tg(slc1a3b:TagRFP-CAAX);Tg(elavl3:H2B-GCaMP6s)* larvae. Of the scyllate heads that became LysoTracker^+^, all contained GCaMP^+^ nuclei (7/7 scyllate heads from 3 fish, Figure 4H). From this data, we conclude that scyllate heads can acidify when holding neuronal cell bodies, but that this phenomenon is rare and astroglia do not usually degrade contents in scyllate heads.

### Microglia interact with scyllate heads and remove their contents

Having observed that scyllate heads are infrequently acidic, rarely internalized into the astroglial soma, and usually dissipate in a bursting or collapsing manner, we investigated the fate of scyllate heads and their contents. While we show that scyllate heads contain dying neurons, most developmental efferocytosis is performed by microglia^12,39,41^. By the time of robust scyllate head formation at 60 hpf, only a small number of microglia have infiltrated into the OT^12,13^. Therefore, we sought to determine if and how microglia and radial astroglia interact during clearance of dying cells at this developmental time point.

We first looked at time-lapses in which we had labeled microglia alongside radial astroglia in *Tg(slc1a3b:myrGFP-P2A-H2AmCherry);Tg(mfap4:mTurquoise)* larvae between 55 hpf and 84 hpf. In these videos we observed that scyllate heads would often dissipate after microglia had been near them (data not shown). Therefore, we hypothesized that microglia directly interact with scyllate heads and cause their dissipation.

Microglia are extremely motile when responding to debris, so to capture microglial dynamics, we imaged *Tg(slc1a3b:myrGFP-P2A-H2AmCherry);Tg(mfap4:mTurquoise)* larvae from 77 to 79 hpf in 110 second time-steps. In our time-lapses, we observed multiple instances of microglia interacting with scyllate heads. In the representative time-lapse shown, we observed a microglial process contact a scyllate head, then enter and form its own engulfment compartment inside the scyllate head (Figure 5A). The microglial process then retracted and the scyllate head rapidly dissipated (Figure 5A). This led us to hypothesize that microglia remove contents from scyllate heads, leading to scyllate head dissipation.

**Figure 5.**
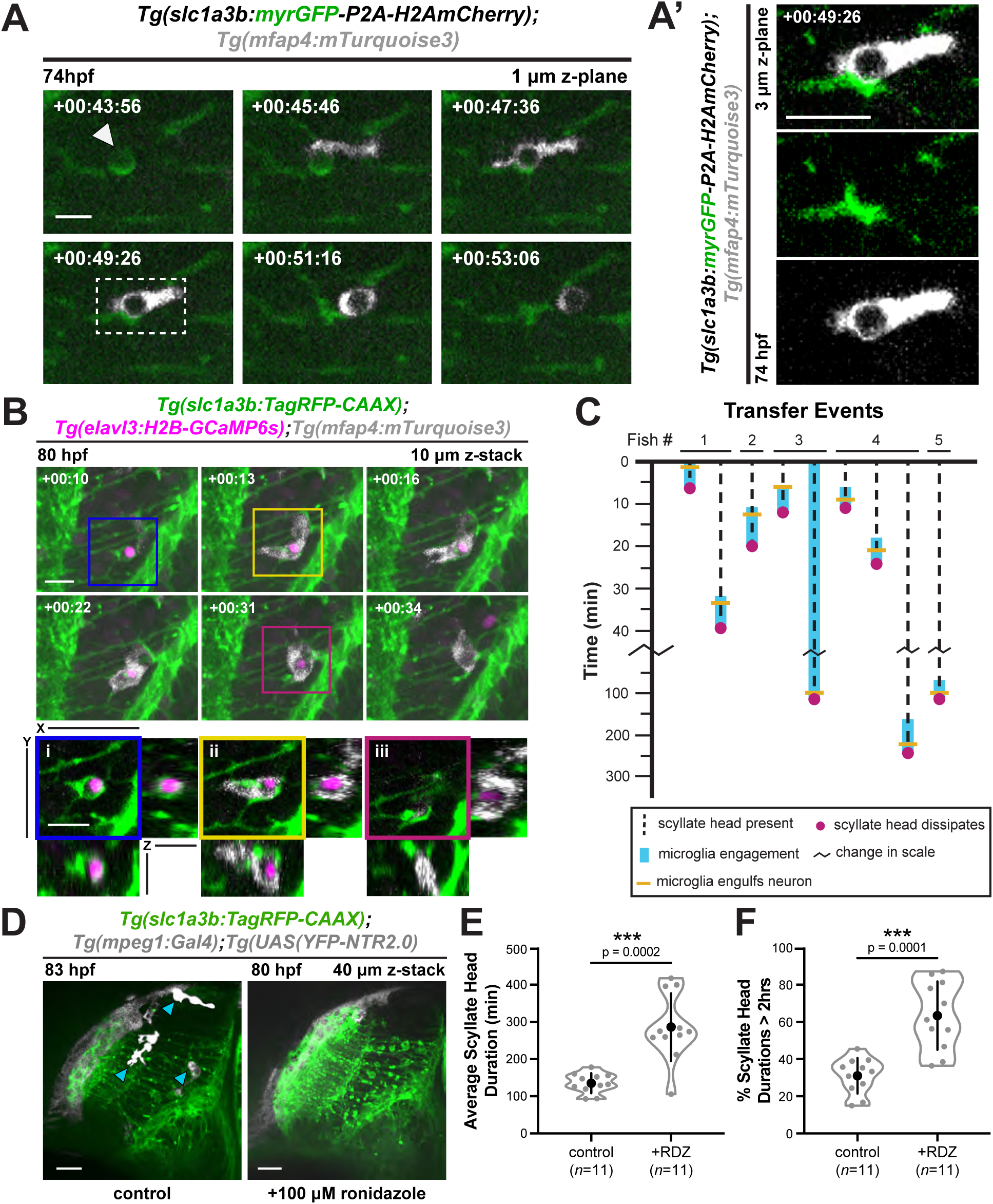
Microglia interact with scyllate heads to remove contents for terminal degradation. A) Single z-plane still images from a time-lapse of a microglial process (white) infiltrating a scyllate head (green, arrowhead) in a *Tg(slc1a3b:TagRFP-CAAX);Tg(elavl3:H2B-GCaMP6s);Tg(mfap4:mTurquoise)* larva at 74 hpf. Dotted lines indicate inset area in A’. A’) Insets from A showing separate channels. B) 10 µm projection images from a time-lapse of a scyllate head-microglial transfer event in a *Tg(slc1a3b:TagRFP-CAAX);Tg(elavl3:H2B-GCaMP6s);Tg(mfap4:mTurquoise)* larva. Colored boxes correspond to insets below. Insets show isometric views of: i) scyllate head holding neuron. ii) microglia infiltrating scyllate head and surrounding neuron. iii) microglia removing neuron from dissipating scyllate head. C) Temporal traces of representative transfer events of neuronal nuclei from scyllate heads to microglia, from scyllate head formation to dissipation. Traces measured from 2-hour timelapses of *Tg(slc1a3b:myrGFP-P2A-H2AmCherry);Tg(elavl3:H2B-GcaMP6s);Tg(mfap4:mTurquoise)* larvae between 76 and 81 hpf. *n*=9 transfer events from 5 fish. D) 40 µm projection images of *Tg(slc1a3b:TagRFP-CAAX);Tg(mpeg1:Gal4);Tg(UAS:YFP-NTR2.0)* larvae in control and after 24 hr 100 µM ronidazole (RDZ) treatment. *Mpeg1+* microglia (white) are indicated with cyan arrowheads. E) Quantification of scyllate head duration averages from time-lapses of *Tg(slc1a3b:TagRFP-CAAX);Tg(mpeg1:Gal4);Tg(UAS:YFP-NTR2.0)* larvae from 75 to 87 hpf in controls and after microglial depletion via RDZ. Overall averages=135±27 minutes in control larvae and 286±91 minutes in microglia-depleted larvae (p=0.0002, *n*=11 fish per condition). F) Quantification of percent of scyllate heads with durations over 2 hours from time-lapses of *Tg(slc1a3b:TagRFP-CAAX);Tg(mpeg1:Gal4);Tg(UAS:YFP-NTR2.0)* larvae from 75 to 87 hpf in controls and after microglial depletion with 24 hr 100 µM RDZ treatment. Averages= 31.1±9.6% in control larvae and 63.3±18.4% in microglia-depleted larvae (p < 0.0001, *n*=11 fish per condition). Time stated in hh:mm:ss (A) or hh:mm (B) after start of imaging. Scale bars, 10 μm (A-B), 20 µm (D).

To directly show that microglia removed neuronal debris from scyllate heads, we used *Tg(slc1a3b:TagRFP-CAAX);Tg(elavl3:H2B-GCaMP6s);Tg(mfap4:mTurquoise)* larvae to label astroglia, microglia, and neurons in the same fish. We imaged these larvae between 73 to 78 hpf in 2-hour time-lapses with 90 to 120 second time-steps. In these time-lapses, we observed instances of neuron transfer from scyllate heads to microglia, which we call transfer events. In the representative time-lapse shown in Figure 5B, we observed a microglial cell contact a scyllate head that contained a GCaMP^+^ neuron. The microglia then infiltrated the scyllate head and surrounded the neuron with its own membrane processes. The microglia then moved away from the scyllate head, taking the neuron with it. During this transfer event, the scyllate head rapidly dissipated. Another representative transfer event shown in Video S5 follows a similar process. On average, we observed 3.6 transfer events per fish during our 2-hour time-lapses (*n*=5 fish). Our observations of glial-glial debris transfer events are supported by a recent study which found that dead cells in the zebrafish retina undergoing phagocytosis by Müller glia were often transferred to microglia for terminal degradation^39^.

To further characterize these transfer events, we tracked individual scyllate heads contacted by microglia throughout our time-lapses (Figure 5C). The time from scyllate head formation to the point when a microglial cell first contacted the scyllate head, which we called microglia engagement, varied greatly, ranging from 0 minutes to 146 minutes (median=21 minutes, average=35±39 minutes, *n*=18 scyllate heads from 5 fish). In most cases, the time from microglial engagement to neuron engulfment and subsequent scyllate head dissipation was short (median=6 minutes, average=14±19 minutes), with a few notable exceptions (see Figure 5C, trace 5). Of the 19 scyllate heads we observed dissipate during our time-lapses, 18 dissipated after a microglial cell infiltrated and removed the neuronal contents (*n*=5 fish). Taken together, we conclude that most scyllate heads dissipate after transferring their debris to microglia.

We further investigated the manner of scyllate head dissipation after a transfer event. In our imaging, microglial compartments appeared to form inside of scyllate heads and only surrounded the contained neuronal debris rather than forming around the entire scyllate head (Figure 5A’, 5Bii). We did not observe scyllate head debris within the microglial compartment containing the transferred neuron after removal (Figure 5Biii). Additionally, we note that in our time-lapses of mosaically labeled astroglia in which we captured scyllate heads dissipating, the scyllate head appeared to burst and be reabsorbed into the basal process (Figure 1F, Video S2). If the scyllate head in this time-lapse were pruned by a microglial cell, we would expect to see scyllate head debris detach and move away from the basal process, which we did not observe. Our data is consistent with the hypothesis that microglia infiltrate scyllate heads to remove their contents and do not prune the entire scyllate head structure.

Interestingly, in our 6 to 12 hour time-lapses of 55 to 84 hpf *Tg(slc1a3b:myrGFP-P2A-H2AmCherry);Tg(mfap4:mTurquoise)* larvae treated with LysoTracker, we did not observe any transfer events between microglia and LysoTracker^+^ scyllate heads. This is consistent with our observations that acidified scyllate heads are longer-lasting and dissipate less often during our time-lapses (Figure 4G-H). However, the 5-minute time-step of these time-lapses and rarity of scyllate head acidification limited our ability to assay these events. Further work is needed to determine if microglia can interact with acidified scyllate heads. Taken together, we conclude that the majority of scyllate heads are contacted by microglia, leading to infiltration of the scyllate head and removal of neuronal debris by microglia for terminal degradation, resulting in dissipation of the scyllate head. This data reveals cooperation between OT radial astroglia and microglia in the clearance of dying neurons during development.

### Depletion of microglia leads to longer lasting scyllate heads

We hypothesized that microglia remove contents from astroglia that would otherwise not be effectively degraded, and that transfer events are important for scyllate head dissipation and dead cell clearance. From this hypothesis, we predicted that loss of microglia would lead to longer lasting scyllate heads. Therefore, we sought to deplete microglia and observe the effects on scyllate head duration.

To deplete microglia, we utilized the nitroreductase (NTR)-ronidazole (RDZ) ablation system, where NTR^+^ cells die when treated with RDZ^39^. We treated *Tg(slc1a3b:TagRFP-CAAX);Tg(mpeg1:Gal4);Tg(UAS:YFP-NTR2.0)* larvae with 100 μM RDZ for 24 hours prior to and throughout imaging, leading to depletion of *mpeg1^+^* microglia (Figure 5D). We then imaged RDZ-treated and untreated larvae from 75 to 87 hpf following microglial depletion and tracked individual scyllate heads to determine their duration. In microglia-depleted larvae, average scyllate head duration was significantly longer than that of untreated larvae (286±91 minutes in RDZ treated larvae vs 135±27 minutes in control larvae, *n*=11 fish per condition, p=0.0002; Figure 5E). Additionally, the distribution of durations was significantly different between the control and microglia-depleted conditions (Figure S3A). The proportion of scyllate heads that lasted longer than 2 hours (the approximate median duration we previously determined, see Figure 1E) in microglia-depleted larvae was double that of untreated larvae (63.3±18.4% in RDZ-treated larvae vs 31.1±9.6% in control larvae, *n*=11 fish per condition, p=0.0001; Figure 5F). Taken together, our data demonstrates that microglia are important for scyllate head dissipation and when they are depleted, scyllate heads continue to hold onto their contents. This is consistent with our hypothesis that microglia and astroglia cooperate to clear neuronal debris during OT development.

## Discussion

In this study, using *in vivo,* time-lapse imaging, we show that radial astroglia in the developing zebrafish OT respond *en masse* to dying neurons during a wave of programmed neuronal cell death by surrounding and holding neuronal cell bodies in structures we call scyllate heads, revealing a previously undescribed developmental phenomenon in the OT. We also show that scyllate heads are commonly contacted by microglia, leading to infiltration into the compartment by microglial processes and subsequent transfer of scyllate head contents to the microglial cell. Our research reveals that astroglia actively participate and coordinate with microglia during the response to OT developmental neuronal cell death.

### OT Radial Astroglia as First-Responders

Previous studies describe astrocytes and astrocyte-like cells as non-professional phagocytes which, in the absence of microglia, will increase their phagocytic capacity to partially, though less effectively, compensate^7,8,21^. Under homeostasis, mouse astrocytes actively prune synapses^43^ and respond to neuronal cell death by engulfing neurite debris^21^. However, whether astroglial cells participate in whole cell clearance under homeostatic conditions is unclear^21,45^. In our studies, we observed OT astroglia identify and surround neuronal cell bodies in scyllate heads during physiological development, even while microglia were active. Occasionally, we observed scyllate heads acidify or be internalized into the astroglial soma, sometimes accompanied by the debris signal breaking apart or being quenched, revealing that OT radial astroglia can phagocytose neuronal cell bodies. However, despite responding to the majority of dying neurons, most scyllate heads do not terminally degrade their contents, leading us to wonder why there is such a robust astroglial response to neuronal cell death.

Most studies on efferocytosis focus on the cells which terminally degrade apoptotic cells, which may have limited our identification of the contributions by non-professional phagocytes to homeostatic cell clearance. For example, a previous study on non-professional phagocytosis briefly described, but did not show, mouse BHK cells *in vitro* surrounding apoptotic cells by forming “a long process” which “mov[ed the apoptotic cell] about”, a description which resembles a scyllate head^7^. However, this study did not consider the BHK cells as ‘phagocytic’ until internalization many hours later and thus did not elaborate on this pre-phagocytic behavior^7^. By expanding our view of what constitutes participation in cell clearance, our study and others have begun to paint non-professional phagocytes as ‘first responders’ to cell death, rather than bystanders that only react when professional phagocytes fail.

In our study, we propose that OT radial astroglia form scyllate heads as an intermediary compartment to temporarily hold neurons until they are cleared by microglia. The purpose of an intermediary compartment, especially one as dynamic and likely metabolically costly as a scyllate head, remains unanswered. We hypothesize that scyllate heads are so numerous and long-lasting because, at this point in OT development, the microglia present are not able to effectively clear the large number of dying cells. Because radial astroglial membranes span the OT, they are in close proximity to neurons and thus are well-poised to quickly respond to neuronal cell death. We observed a decrease in scyllate head formation after 4 dpf, which may be because there are more microglia able to efficiently clear the lower number of apoptotic neurons. If dead cells are not cleared efficiently, they begin to secrete inflammatory and cytotoxic cues^46,47^. Therefore, we hypothesize that scyllate heads form to sequester dying neurons, keeping the dead cells from harming nearby healthy neurons until microglia are able to remove the debris for degradation.

In addition to debris sequestering, there could be other, not necessarily mutually exclusive, benefits to astroglial involvement in cell clearance. For example, in mice, mammary epithelial cells acting as non-professional phagocytes secrete the anti-inflammatory cytokine TGF-β after engulfing apoptotic cells^6^. It is possible that, during this time of high death, the containment of apoptotic cells within scyllate heads could stimulate a similar release of protective cues by astroglia. Another possibility is that scyllate heads secrete signals which help guide invading microglia into the CNS, similar to how astrocytes in mouse models of Alzheimer’s Disease secrete C3 and influence microglial activity^44,45^. Scyllate heads form around dying cells concurrent to an increase in PCD, and microglial infiltration into the OT is dependent on the amount of cell death at this time-point^12,14^, so scyllate heads may also contribute to microglial invasion. However, we do not currently have the tools to specifically perturb scyllate head formation or function without completely eliminating radial astroglia and/or disrupting overall OT function, and thus more work is needed to investigate what may happen in the OT in the absence of scyllate heads.

### Transfer of Cellular Debris from Radial Astroglia to Microglia

Recently, investigations into interactions between astroglia and microglia have revealed intimate communication between these two cell types. In the developing brains of both mice and rhesus monkeys, microglia closely contact and survey radial glial projections, though the purpose of the interaction is unclear from these studies^48,49^. In the mouse hypothalamus, astrocytes and microglia coordinate during clearance of ablated neurons by partitioning the types of neuronal debris each one clears, with astroglia engulfing mostly neurite debris^21^.

In our study, we utilized *in vivo,* time-lapse imaging in zebrafish to interrogate interactions between OT radial astroglial scyllate heads and microglia. We show that OT radial astroglia and microglia coordinate during developmental clearance of apoptotic cells to transfer neuronal debris from scyllate heads to microglial phagosomes. To our knowledge, only one other study has identified a similar phenomenon of direct debris transfer between non-professional and professional phagocytes, and only very recently^39^. In this study, Morales et al.^39^ observed transfer of apoptotic debris from Müller glia to microglia during a wave of apoptosis in the developing zebrafish retina. In our work, we show that transfer events occur in the zebrafish OT and are not unique to Müller glia-microglia interactions. In addition, our imaging studies further elucidate the cellular interactions involved in transfer events, such as the initial contact between the scyllate heads and microglia and subsequent ‘opening’ of scyllate heads, as well as the specific type of debris involved (e.g. neuronal cell bodies). Our high-speed imaging reveals that scyllate heads are infiltrated by microglia rather than being pruned along with their contents as reported by Morales et al.^39^, indicating that microglia-scyllate head transfer events are highly coordinated and may require complex cell-cell communication. Considering similar phenomena occur in these two different contexts, the brain and retina, we suspect that transfer events between professional and non-professional phagocytes may be a common occurrence during cell clearance.

### A Model for Professional and Non-professional Phagocyte Cooperation

Based on our observations, we propose a model of *in vivo* efferocytosis during OT development in which radial astroglia initially surround apoptotic neuronal targets in a temporary membranous holding compartment, which we call a scyllate head. Occasionally, scyllate heads can become degradative, leading to terminal efferocytosis of neuronal cells by radial astroglia. However, in most cases, microglia contact, invade, and remove contents from scyllate heads for terminal degradation instead. Combined with observations in other non-professional phagocytes like zebrafish Müller glia^39^ and mouse BHK cells^7^, we hypothesize that this process of initial compartmentalization of apoptotic cells by non-professional phagocytes and later transfer to professional phagocytes could be generalized to other biological contexts.

To better understand the contribution of radial astroglia to cell clearance, future studies are needed to determine the cellular mechanisms involved in scyllate head formation and interactions with microglia and to create tools to specifically disrupt them. Known astroglial phagocytic receptors that may be involved in scyllate head target recognition include *mertk/axl* and *megf10,* also called Draper in *Drosophila*^2,20,21,39,43^. It is possible that, as intermediate compartments, scyllate heads could utilize different receptors and cues than astroglial phagosomes that engulf synapses and neurites. Our data shows that most scyllate head contents are at a stage in which they have exposed phosphatidylserine (secA5-YFP^+^) but not undergone chromatin condensation (AO^+^), which may mean that scyllate heads respond to early signals from apoptotic or even pre-apoptotic cells. Even more unclear are the potential cell-cell communication pathways between scyllate heads and microglial processes, including how microglia find apoptotic debris within a scyllate head and how microglia enter scyllate heads without engulfing them. Finally, many studies have sought to identify what cues cause non-professional phagocytes to actively engulf and degrade debris. Parnaik et al.^7^ suggests this switch is time-dependent, but whether it is due to a change in the apoptotic cell or the phagocytic ability of the non-professional phagocyte itself is debated^6,7^. Our findings suggest that scyllate head acidification is not wholly dependent on scyllate head duration, however, we observed terminal degradation by radial astroglia too rarely to make a conclusion. We note that, despite experimental caveats, LysoTracker^+^ scyllate heads did not dissipate due to microglial interaction. This could indicate an intrinsic change from a scyllate head acting as an intermediate holding compartment and a scyllate head that has acidified and potentially becoming degradative that precludes microglial interaction. Our work further invites investigation into the shift in non-professional phagocytes from first responders to active phagocytes under certain conditions and what cues may cause this change.

### Resource Availability

#### Lead Contact

Requests for further information and resources should be directed to and will be fulfilled by the lead contact, Sarah Kucenas.

#### Materials Availability

All materials, reagents, and transgenic lines generated in this article are available upon request to the lead contact. Detailed protocols for experiments reported in this article are available upon request to the lead contact.

#### Data Availability

Quantification data, original images, and analyzed images are available upon request to the lead contact.

## Acknowledgements

We thank members of the Kucenas lab, past and present, for valuable discussions and training, and especially Lori Tocke for zebrafish care. We thank Drs. Noelle Dwyer, Xiaowei Lu, Manoj Patel, and Ann E. Sutherland for guidance and training. We also thank Dr. Kelly Monk for the *Tg(slc1a3b:myrGFP-P2A-H2AmCherry)* line and related constructs, Dr. Laura Fontenas for the pMe-myrEGFP construct, and Dr. David Schoppik for the *lakritz* mutant and *Tg(ath5:mRFP)* lines. This work was funded by the NIH/NEI (F31EY035961, HMB), the Robert R. Wagner Fellowship Fund (HMB), the Owens Family Foundation (SK), and the NIH/NINDS (R01NS107525, SK).

## Author Contributions

Conceptualization, H.M.B. and S.K.; Methodology, H.M.B. and S.K.; Investigation, H.M.B., C.G.R., and S.K.; Writing – Original Draft, H.M.B and S.K.; Writing – Review & Editing, H.M.B. and S.K.; Formal analysis, H.M.B., C.G.R., and Z.C.; Transgenic line generation, H.M.B., R.I.B., Z.C., and I.W.

## STAR Methods

### Fish husbandry

All animal studies were approved by the University of Virginia Institutional Animal Care and Use Committee (IACUC). Animals were housed at 28°C at a density of 8 to 10 fish/liter and exposed to a 14/10 hour light-dark cycle. Previously published and newly generated zebrafish strains used in this study are included in Table 1. Embryos were produced by pairwise natural matings and raised in egg water at 28.5°C. Larvae were staged according to hours post fertilization (hpf) and screened for desired transgenic fluorescence prior to use. Sex was not considered as a variable for experiments because sex is indeterminate in zebrafish prior to 25 dpf.

### Generation of transgenic lines

Lines generated in this study include *Tg(slc1a3b:mCerulean)^uva84^, Tg(elavl3:H2B-GCaMP6s)^uva98^,* and *Tg(slc1a3b:myrEGFP)^uva74^*. Transgenic constructs were created using the Tol2kit Gateway-based cloning system^46^. The slc1a3b-p5e plasmid was provided by Kelly Monk^19^. The myrEFGP-pMe plasmid was provided by Laura Fontenas and produced from the *slc1a3b:myrGFP-P2A-H2AmCherry* plasmid provided by Kelly Monk^19^. AB* embryos were injected with 20 ng/μL plasmid DNA and 100 ng/μL transposase mRNA. Embryos were screened for expression and positive embryos were grown to adulthood. Founders were determined by pairwise individual matings with AB* adults and progeny were screened for desired expression. To generate a stable line, outcrosses were performed for two generations.

### Genotyping

Genotype of imaged embryos was determined post-imaging for relevant experiments. After imaging, larvae were individually removed from imaging dishes. DNA was extracted using HotSHOT lysis solutions and protocol^48^. PCR amplification was performed using GoTaq Green Master Mix (Fisher) with corresponding primers found in the Key Resources Table. Genotype of *lakritz* mutants was determined by performing a restriction digest on the PCR product with AcuI restriction enzyme (New England Biolabs).

### Mosaic labeling of cells

One-cell embryos were injected with *slc1a3b:myrGFP-P2A-H2AmCherry* plasmid DNA (provided by Kelly Monk) as described in the “Generation of transgenic lines” section. Larvae were screened for desired fluorescence and those with labeled OT radial glia were subsequently imaged.

### In vivo imaging

Embryos were treated with 0.003% PTU at 24 hpf to reduce pigment formation and screened for transgenic fluorescence prior to imaging. For time points before 56 hpf, embryos were manually dechorionated. Larvae were immobilized prior to imaging using diluted Tricaine (MS-222; Syndel/Western Chemical, ordered from The Pond Outlet; concentration varied by age). Larvae were then mounted on their sides in 0.8% low gelling point agarose in 4-well glass bottom 35 mm Petri dishes (Fisher, Greiner Bio-One) and covered with diluted Tricaine (concentration varied by age), unless otherwise noted. Imaging was performed on either Andor CSU-W1 Spinning Disc Confocal Microscope or Andor Dragonfly 200 Spinning Disc Confocal Microscope with iXon Ultra camera with 40x/1.10 objective (Andor/Oxford Instruments). Images were illuminated with 445 nm (mTurquoise, mCerulean), 488 nm (GFP, GCaMP6s, YFP, AO), 561 nm (mCherry, TagRFP), and/or 647 nm (LysoTracker) lasers. Time-lapse length and time-step varied according to the feasibility and needs of the experiment and is stated if relevant. Images were acquired as 40 μm z-series with 1 μm step size unless otherwise stated. For some representative images, imaging was performed using frame averaging of 4 (Figure 3, Video S4).

### Microglial depletion

To ablate microglia via nitroreductase-ronidazole (NTR-RDZ) treatment, *Tg(mpeg1:Gal4);Tg(UAS:YFP-NTR2.0)* larvae were used. Expression of the *YFP-NTR2.0* was confirmed prior to treatment. Egg water was replaced with 100 μM RDZ (Sigma) dissolved in PTU approximately 12 or 24 hours prior to imaging. Control siblings were kept in PTU only. When it was not possible to screen larvae prior to RDZ treatment, positive larvae were identified on the confocal microscope prior to imaging by the presence of low *mpeg1*-expressing neurons in the spinal cord. Treated larvae remained in 100 μM RDZ throughout imaging.

### Acridine Orange incorporation

Acridine Orange (AO) treatment was performed as described in Lyons et al., 2005^49^. Briefly, larvae were immersed in a solution of 10 μg/mL AO (Santa Cruz Biotechnology) in the dark at 28.5°C for 20 minutes. Larvae were washed in egg water three times. Larvae were then immobilized and mounted for imaging as previously described. If necessary, embryos were dechorionated prior to treatment.

### LysoTracker treatment

LysoTracker treatment was performed as described in Zhu et al. 2019^12^. Briefly, larvae were immersed in 10 μM LysoTracker Deep Red (Invitrogen) in the dark at 25 °C for 1 hour. The larvae were rinsed with egg water three times and then immobilized and mounted for imaging as previously described. If necessary, embryos were dechorionated prior to treatment.

### Image analysis

Time-lapse images were analyzed using Imaris software (Version 10.2.1). To standardize analysis, only astroglia in the dorsal anterior side of the OT (i.e. those projecting through the SPV and neuropil, see Figure S1A) were considered for analysis. Scyllate heads were identified by assessing individual z-planes for a spherical or hemi-spherical shape and evaluating nearby time points for characteristic formation and movement. In *Tg(slc1a3b:myrGFP-P2A-H2AmCherry)* larvae, spherical shapes containing *mCherry^+^* nuclei were not considered scyllate heads regardless of position to avoid confusion with astroglial cell bodies. For Figure 1C, we examined frames at predetermined time-points from time-lapses and counted the number of scyllate heads visible in that frame. We examined 2-3 time-points per time-lapse, based on which time-points were covered by the time-lapse.

The “spots’’ tool was used to track individual scyllate heads manually. For analysis of individual scyllate heads, spot tracks were numbered in order of appearance first and disappearance second. Similar procedures were followed for tracking secA5-YFP^+^ neurons (Figure 3B-C, 4B).

Scyllate head size (Figure 1D) was measured in Imaris using the distance tool. Scyllate heads were measured at the time-point and single z-plane at which they appeared largest by eye.

Scyllate head duration (Figure 1E, 4G, 5E-F) was calculated as the difference between the first frame and last frame that the scyllate head could be identified multiplied by the time-step between frames. Scyllate heads present at the beginning of the time-lapse were counted as appearing at the first frame and scyllate heads present at the end of the time-lapse were counted as dissipating at the last frame. To determine the time to scyllate head engagement (Figure 3F), we only considered neurons which became secA5-YFP^+^ during the time-lapse (i.e. did not begin secA5-YFP^+^) and were at some point surrounded by a scyllate head. The time to engagement was calculated as the difference between the frame in which the neuron became recognizably secA5-YFP^+^ and the frame in which a cup-like astroglial projection contacted the neuron, multiplied by the time-step. To determine time to acidification (Figure 4F), we only considered scyllate heads which became LysoTracker^+^ during the timelapse. The time to acidification was calculated as the difference between the frame at which the scyllate head formed and the frame at which the scyllate head contained LysoTracker^+^ signal discernable above background, multiplied by the time-step.

When determining the proportion of scyllate heads that became GCamp^+^ (Figure 3B), AO^+^ (Figure S2E), and LysoTracker^+^ (Figure 4D), scyllate heads were tracked for their entire duration and examined for relevant characteristics throughout the time-lapse. Scyllate head contents were considered positive for a marker if fluorescence was discernibly above background. When determining the proportion of secA5-YFP^+^ neurons that were engaged by scyllate heads (Figure 3D) or internalized (Figure 4B), secA5-YFP^+^ neuron cell bodies were tracked throughout the entire duration of discernable fluorescence. They were considered engaged by a scyllate head when they were contacted by cup-like astroglial membrane that would surround them later in the time-lapse. Those contained in scyllate heads were considered internalized if they were moved to an astroglial cell body and the scyllate head no longer formed a clear boundary around the secA5-YFP^+^ debris.

Brightness and contrast of images were adjusted uniformly throughout all images, with each channel adjusted individually, to improve image clarity. Original, unprocessed images and images with Imaris analysis are available upon request.

### Statistical methods

Prism or R software was used to perform statistical analyses of quantifications and create graphs. Prism and R analysis documents and raw quantification data are available upon request. Graph sizing and aesthetic presentation were adjusted in Illustrator without any changes made to the data shown. All individual data points used for graphical generation are shown. Error bars indicate standard deviation unless otherwise stated. Results are presented in mean±standard deviation format unless otherwise stated. Sample size (*n*) is reported in the results and in the figure and figure legend where applicable. What *n* represents is specified in the results.

For Figure 2F, to elucidate the margin of error caused by the chosen time-step, data is portrayed as a histogram with 10-minute intervals such that events with a time to engagement of 0 minutes were in the 0-9 minute bin. Unpaired t-tests with Welch’s correction were performed for data shown in Figure 5E-F. In Figure 5E-F, individual data points shown represent the average scyllate head duration of each fish, with the overall average per condition being the average of these averages. For further transparency, Figure S3A-A’ shows the individual scyllate head durations for all fish in each condition and for each fish individually. A Kolmogorov-Smirnov test was performed for data shown in Figure S3A.

## Supplemental Item Titles and Legends

**Document S1.** Table S1 and Supplemental References

**Table S1. Transgenic and mutant fish lines used in this study, related to STAR Methods.**

**Video S1. OT astroglia form scyllate heads during a critical developmental period, related to** Figure 1. Time-lapse of a *Tg(slc1a3b:myrGFP-P2A-H2AmCherry)* larva, 40 μm projection. Astroglial nuclei (magenta) and membranes (green) are labeled. Scyllate heads are indicated with arrowheads at their first appearance. Time indicated in hh:mm:ss after start of imaging at 72 hpf. Scale bar, 10 μm.

**Video S2. Single OT astroglia forming scyllate head, related to** Figure 1. Time-lapse of a single radial astroglia labeled with *slc1a3b:myrGFP-P2A-H2A-mCherry*. The astroglia (green membranes, magenta nucleus) forms a scyllate head from its basal projection, indicated by white arrowhead. The scyllate head later dissipates, indicated by arrowhead outline. Time indicated in hh:mm:ss after start of imaging at 57 hpf. Scale bar, 10 µm.

**Video S3. *Lakritz* mutant larvae lack RGC axons but form scyllate heads during development, related to** Figure 2. Time-lapses of *Tg(slc1a3b:myrGFP-P2A-H2AmCherry);Tg(ath5:mRFP)*;*lakritz (lak)* heterozygous (left) and homozygous (right) larvae. RGC axons (magenta) can be seen infiltrating the *lak/+* OT but not the sibling *lak/lak* OT. Scyllate heads (green) are indicated by arrowheads at formation. Time indicated in hh:mm:ss after start of imaging at 51 hpf. Scale bar, 20 µm.

**Video S4. Scyllate heads surround dying neurons, related to Figure 3**. Time-lapse of a *Tg(slc1a3b:TagRFP-CAAX);Et(s1101:Gal4);Tg(UAS:secA5-YFP)* larva, 40 µm projection. Dying neuron cell bodies (magenta) surrounded by scyllate heads (green) are indicated with arrowheads at time of scyllate head engagement. Time indicated in hh:mm:ss after start of imaging at 54 hpf. Scale bar, 10 µm.

**Video S5. Microglia remove neuronal cell bodies from scyllate heads.** Time-lapse of a *Tg(slc1a3b:TagRFP-CAAX);Tg(elavl3-H2b-GCaMP6s);Tg(mfap4:mTurquoise)* larva, 10 µm projection. A scyllate head (arrowhead, green) forms around a neuronal cell body (magenta) and is later infiltrated by a microglial projection (white) which removes the neuron. During this process, the scyllate head dissipates. Arrow indicates the start of microglial engagement. Time indicated in hh:mm:ss after start of imaging at 76 hpf. Scale bar, 5 µm.

## References

1. Hofmann K. The Evolutionary Origins of Programmed Cell Death Signaling. Cold Spring Harb Perspect Biol. 2020 Sep 1;12(9):a036442. doi: 10.1101/cshperspect.a036442. PMID: 31818855; PMCID: PMC7461761.

2. Liu KE, Raymond MH, Ravichandran KS, Kucenas S. Clearing Your Mind: Mechanisms of Debris Clearance After Cell Death During Neural Development. Annu Rev Neurosci. 2022 Jul 8;45:177–198. doi: 10.1146/annurev-neuro-110920-022431. Epub 2022 Feb 28. PMID: 35226828; PMCID: PMC10157384.

3. Boada-Romero E, Martinez J, Heckmann BL, Green DR. The clearance of dead cells by efferocytosis. Nat Rev Mol Cell Biol. 2020 Jul;21(7):398–414. doi: 10.1038/s41580-020-0232-1. Epub 2020 Apr 6. PMID: 32251387; PMCID: PMC7392086.

4. Ge Y, Huang M, Yao YM. Efferocytosis and Its Role in Inflammatory Disorders. Front Cell Dev Biol. 2022 Feb 25;10:839248. doi: 10.3389/fcell.2022.839248. PMID: 35281078; PMCID: PMC8913510.

5. Witting A, Müller P, Herrmann A, Kettenmann H, Nolte C. Phagocytic clearance of apoptotic neurons by Microglia/Brain macrophages in vitro: involvement of lectin-, integrin-, and phosphatidylserine-mediated recognition. J Neurochem. 2000 Sep;75(3):1060–70. doi: 10.1046/j.1471-4159.2000.0751060.x. PMID: 10936187.

6. Monks, J., Rosner, D., Jon Geske, F. et al. Epithelial cells as phagocytes: apoptotic epithelial cells are engulfed by mammary alveolar epithelial cells and repress inflammatory mediator release. Cell Death Differ 12, 107–114 (2005). 10.1038/sj.cdd.4401517

7. Parnaik R, Raff MC, Scholes J. Differences between the clearance of apoptotic cells by professional and non-professional phagocytes. Curr Biol. 2000 Jul 13;10(14):857–60. doi: 10.1016/s0960-9822(00)00598-4. PMID: 10899007.

8. Magnus T, Chan A, Linker RA, Toyka KV, Gold R. Astrocytes are less efficient in the removal of apoptotic lymphocytes than microglia cells: implications for the role of glial cells in the inflamed central nervous system. J Neuropathol Exp Neurol. 2002 Sep;61(9):760–6. doi: 10.1093/jnen/61.9.760. PMID: 12230322.

9. Shklover J, Levy-Adam F, Kurant E. Apoptotic Cell Clearance in Development. Curr Top Dev Biol. 2015;114:297–334. doi: 10.1016/bs.ctdb.2015.07.024. Epub 2015 Sep 9. PMID: 26431572.

10. Niell CM, Smith SJ. Functional imaging reveals rapid development of visual response properties in the zebrafish tectum. Neuron. 2005 Mar 24;45(6):941–51. doi: 10.1016/j.neuron.2005.01.047. PMID: 15797554.

11. Stuermer CA. Retinotopic organization of the developing retinotectal projection in the zebrafish embryo. J Neurosci. 1988 Dec;8(12):4513–30. doi: 10.1523/JNEUROSCI.08-12-04513.1988. PMID: 2848935; PMCID: PMC6569580.

12. Casano AM, Albert M, Peri F. Developmental Apoptosis Mediates Entry and Positioning of Microglia in the Zebrafish Brain. Cell Rep. 2016 Jul 26;16(4):897–906. doi: 10.1016/j.celrep.2016.06.033. Epub 2016 Jul 14. PMID: 27425604.

13. Herbomel P, Thisse B, Thisse C. Zebrafish early macrophages colonize cephalic mesenchyme and developing brain, retina, and epidermis through a M-CSF receptor-dependent invasive process. Dev Biol. 2001 Oct 15;238(2):274–88. doi: 10.1006/dbio.2001.0393. PMID: 11784010.

14. Xu J, Wang T, Wu Y, Jin W, Wen Z. Microglia Colonization of Developing Zebrafish Midbrain Is Promoted by Apoptotic Neuron and Lysophosphatidylcholine. Dev Cell. 2016 Jul 25;38(2):214–22. doi: 10.1016/j.devcel.2016.06.018. Epub 2016 Jul 14. PMID: 27424497.

15. Zhu Y, Crowley SC, Latimer AJ, Lewis GM, Nash R, Kucenas S. Migratory Neural Crest Cells Phagocytose Dead Cells in the Developing Nervous System. Cell. 2019 Sep 19;179(1):74–89.e10. doi: 10.1016/j.cell.2019.08.001. Epub 2019 Sep 5. PMID: 31495570; PMCID: PMC6754278.

16. Czopka T, Monk K, Peri F. Glial Cell Development and Function in the Zebrafish Central Nervous System. Cold Spring Harb Perspect Biol. 2024 Nov 1;16(11):a041350. doi: 10.1101/cshperspect.a041350. PMID: 38692835; PMCID: PMC11529855.

17. Grupp L, Wolburg H, Mack AF. Astroglial structures in the zebrafish brain. J Comp Neurol. 2010 Nov 1;518(21):4277–87. doi: 10.1002/cne.22481. PMID: 20853506.

18. Lyons DA, Talbot WS. Glial cell development and function in zebrafish. Cold Spring Harb Perspect Biol. 2014 Nov 13;7(2):a020586. doi: 10.1101/cshperspect.a020586. PMID: 25395296; PMCID: PMC4315925.

19. Lu, Z., Elliott, M., Chen, Y. et al. Phagocytic activity of neuronal progenitors regulates adult neurogenesis. Nat Cell Biol 13, 1076–1083 (2011). 10.1038/ncb2299

20. Tasdemir-Yilmaz OE, Freeman MR. Astrocytes engage unique molecular programs to engulf pruned neuronal debris from distinct subsets of neurons. Genes Dev. 2014 Jan 1;28(1):20–33. doi: 10.1101/gad.229518.113. Epub 2013 Dec 20. PMID: 24361692; PMCID: PMC3894410.

21. Damisah EC, Hill RA, Rai A, Chen F, Rothlin CV, Ghosh S, Grutzendler J. Astrocytes and microglia play orchestrated roles and respect phagocytic territories during neuronal corpse removal in vivo. Sci Adv. 2020 Jun 26;6(26):eaba3239. doi: 10.1126/sciadv.aba3239. PMID: 32637606; PMCID: PMC7319765.

22. Chen J, Poskanzer KE, Freeman MR, Monk KR. Live-imaging of astrocyte morphogenesis and function in zebrafish neural circuits. Nat Neurosci. 2020 Oct;23(10):1297–1306. doi: 10.1038/s41593-020-0703-x. Epub 2020 Sep 7. PMID: 32895565; PMCID: PMC7530038.

23. Teranikar T, Villarreal C, Salehin N, Ijaseun T, Lim J, Dominguez C, Nguyen V, Cao H, Chuong CJ, Lee J. Scale space detector for analyzing spatiotemporal ventricular contractility and nuclear morphogenesis in zebrafish. iScience. 2022 Aug 5;25(9):104876. doi: 10.1016/j.isci.2022.104876. PMID: 36034231; PMCID: PMC9404658.

24. Kay JN, Finger-Baier KC, Roeser T, Staub W, Baier H. Retinal ganglion cell genesis requires lakritz, a Zebrafish atonal Homolog. Neuron. 2001 Jun;30(3):725–36. doi: 10.1016/s0896-6273(01)00312-9. PMID: 11430806.

25. Kelsh RN, Brand M, Jiang YJ, Heisenberg CP, Lin S, Haffter P, Odenthal J, Mullins MC, van Eeden FJ, Furutani-Seiki M, Granato M, Hammerschmidt M, Kane DA, Warga RM, Beuchle D, Vogelsang L, Nüsslein-Volhard C. Zebrafish pigmentation mutations and the processes of neural crest development. Development. 1996 Dec;123:369–89. doi: 10.1242/dev.123.1.369. PMID: 9007256.

26. Martin A, Babbitt A, Pickens AG, Pickett BE, Hill JT, Suli A. Single-Cell RNA Sequencing Characterizes the Molecular Heterogeneity of the Larval Zebrafish Optic Tectum. Front Mol Neurosci. 2022 Feb 10;15:818007. doi: 10.3389/fnmol.2022.818007. PMID: 35221915; PMCID: PMC8869500.

27. Xiao Y, Petrucco L, Hoodless LJ, Portugues R, Czopka T. Oligodendrocyte precursor cells sculpt the visual system by regulating axonal remodeling. Nat Neurosci. 2022 Mar;25(3):280–284. doi: 10.1038/s41593-022-01023-7. Epub 2022 Mar 3. PMID: 35241802; PMCID: PMC8904260.

28. Clarke, L., Barres, B. Emerging roles of astrocytes in neural circuit development. Nat Rev Neurosci 14, 311–321 (2013). 10.1038/nrn3484

29. Mu Y, Bennett DV, Rubinov M, Narayan S, Yang CT, Tanimoto M, Mensh BD, Looger LL, Ahrens MB. Glia Accumulate Evidence that Actions Are Futile and Suppress Unsuccessful Behavior. Cell. 2019 Jun 27;178(1):27–43.e19. doi: 10.1016/j.cell.2019.05.050. Epub 2019 Jun 20. PMID: 31230713.

30. Chen TW, Wardill TJ, Sun Y, Pulver SR, Renninger SL, Baohan A, Schreiter ER, Kerr RA, Orger MB, Jayaraman V, Looger LL, Svoboda K, Kim DS. Ultrasensitive fluorescent proteins for imaging neuronal activity. Nature. 2013 Jul 18;499(7458):295–300. doi: 10.1038/nature12354. PMID: 23868258; PMCID: PMC3777791.

31. Hou L, Liu K, Li Y, Ma S, Ji X, Liu L. Necrotic pyknosis is a morphologically and biochemically distinct event from apoptotic pyknosis. J Cell Sci. 2016 Aug 15;129(16):3084–90. doi: 10.1242/jcs.184374. Epub 2016 Jun 29. PMID: 27358477.

32. Kerr JF, Wyllie AH, Currie AR. Apoptosis: a basic biological phenomenon with wide-ranging implications in tissue kinetics. Br J Cancer. 1972 Aug;26(4):239–57. doi: 10.1038/bjc.1972.33. PMID: 4561027; PMCID: PMC2008650.

33. Kroemer, G., Galluzzi, L., Vandenabeele, P. et al. Classification of cell death: recommendations of the Nomenclature Committee on Cell Death 2009. Cell Death Differ 16, 3–11 (2009). 10.1038/cdd.2008.150

34. Orrenius S, Zhivotovsky B, Nicotera P. Regulation of cell death: the calcium-apoptosis link. Nat Rev Mol Cell Biol. 2003 Jul;4(7):552–65. doi: 10.1038/nrm1150. PMID: 12838338.

35. Ribble, D., Goldstein, N.B., Norris, D.A. et al. A simple technique for quantifying apoptosis in 96-well plates. BMC Biotechnol 5, 12 (2005). 10.1186/1472-6750-5-12

36. Scott EK, Baier H. The cellular architecture of the larval zebrafish tectum, as revealed by gal4 enhancer trap lines. Front Neural Circuits. 2009 Oct 9;3:13. doi: 10.3389/neuro.04.013.2009. PMID: 19862330; PMCID: PMC2763897.

37. Szobota S, Gorostiza P, Del Bene F, Wyart C, Fortin DL, Kolstad KD, Tulyathan O, Volgraf M, Numano R, Aaron HL, Scott EK, Kramer RH, Flannery J, Baier H, Trauner D, Isacoff EY. Remote control of neuronal activity with a light-gated glutamate receptor. Neuron. 2007 May 24;54(4):535–45. doi: 10.1016/j.neuron.2007.05.010. PMID: 17521567.

38. van Ham TJ, Mapes J, Kokel D, Peterson RT. Live imaging of apoptotic cells in zebrafish. FASEB J. 2010 Nov;24(11):4336–42. doi: 10.1096/fj.10-161018. Epub 2010 Jul 2. PMID: 20601526; PMCID: PMC2974414.

39. Morales M, Findley AP, Mitchell DM. Intercellular contact and cargo transfer between Müller glia and to microglia precede apoptotic cell clearance in the developing retina. Development. 2024 Jan 1;151(1):dev202407. doi: 10.1242/dev.202407. Epub 2024 Jan 4. PMID: 38174987; PMCID: PMC10820749.

40. Moss JJ, Hammond CL, Lane JD. Zebrafish as a model to study autophagy and its role in skeletal development and disease. Histochem Cell Biol. 2020 Nov;154(5):549–564. doi: 10.1007/s00418-020-01917-2. Epub 2020 Sep 11. PMID: 32915267; PMCID: PMC7609422.

41. Mazaheri, F., Breus, O., Durdu, S. et al. Distinct roles for BAI1 and TIM-4 in the engulfment of dying neurons by microglia. Nat Commun 5, 4046 (2014). 10.1038/ncomms5046

42. Curado S, Stainier DY, Anderson RM. Nitroreductase-mediated cell/tissue ablation in zebrafish: a spatially and temporally controlled ablation method with applications in developmental and regeneration studies. Nat Protoc. 2008;3(6):948–54. doi: 10.1038/nprot.2008.58. PMID: 18536643; PMCID: PMC2705989.

43. Chung, WS., Clarke, L., Wang, G. et al. Astrocytes mediate synapse elimination through MEGF10 and MERTK pathways. Nature 504, 394–400 (2013). 10.1038/nature12776

44. Lee SY, Chung WS. The roles of astrocytic phagocytosis in maintaining homeostasis of brains. J Pharmacol Sci. 2021 Mar;145(3):223–227. doi: 10.1016/j.jphs.2020.12.007. Epub 2020 Dec 29. PMID: 33602502.

45. Lian H, Litvinchuk A, Chiang AC, Aithmitti N, Jankowsky JL, Zheng H. Astrocyte-Microglia Cross Talk through Complement Activation Modulates Amyloid Pathology in Mouse Models of Alzheimer’s Disease. J Neurosci. 2016 Jan 13;36(2):577–89. doi: 10.1523/JNEUROSCI.2117-15.2016. PMID: 26758846; PMCID: PMC4710776.

46. Nagata S. Autoimmune diseases caused by defects in clearing dead cells and nuclei expelled from erythroid precursors. Immunol Rev. 2007 Dec;220:237–50. doi: 10.1111/j.1600-065X.2007.00571.x. PMID: 17979851.

47. Ren Y, Savill J. Apoptosis: the importance of being eaten. Cell Death Differ. 1998 Jul;5(7):563–8. doi: 10.1038/sj.cdd.4400407. PMID: 10200510.

48. Barger N, Keiter J, Kreutz A, Krishnamurthy A, Weidenthaler C, Martínez-Cerdeño V, Tarantal AF, Noctor SC. Microglia: An Intrinsic Component of the Proliferative Zones in the Fetal Rhesus Monkey (Macaca mulatta) Cerebral Cortex. Cereb Cortex. 2019 Jul 5;29(7):2782–2796. doi: 10.1093/cercor/bhy145. PMID: 29992243; PMCID: PMC6611465.

49. Rosin JM, Marsters CM, Malik F, Far R, Adnani L, Schuurmans C, Pittman QJ, Kurrasch DM. Embryonic Microglia Interact with Hypothalamic Radial Glia during Development and Upregulate the TAM Receptors MERTK and AXL following an Insult. Cell Rep. 2021 Jan 5;34(1):108587. doi: 10.1016/j.celrep.2020.108587. PMID: 33406432.

50. Kwan KM, Fujimoto E, Grabher C, Mangum BD, Hardy ME, Campbell DS, Parant JM, Yost HJ, Kanki JP, Chien CB. The Tol2kit: a multisite gateway-based construction kit for Tol2 transposon transgenesis constructs. Dev Dyn. 2007 Nov;236(11):3088–99. doi: 10.1002/dvdy.21343. PMID: 17937395.

51. Meeker ND, Hutchinson SA, Ho L, Trede NS. Method for isolation of PCR-ready genomic DNA from zebrafish tissues. Biotechniques. 2007 Nov;43(5):610, 612, 614. doi: 10.2144/000112619. PMID: 18072590.

52. Lyons DA, Pogoda HM, Voas MG, Woods IG, Diamond B, Nix R, Arana N, Jacobs J, Talbot WS. erbb3 and erbb2 are essential for schwann cell migration and myelination in zebrafish. Curr Biol. 2005 Mar 29;15(6):513–24. doi: 10.1016/j.cub.2005.02.030. PMID: 15797019.

53. Brown RI, Barber HM, Kucenas S. Satellite glial cell manipulation prior to axotomy enhances developing dorsal root ganglion central branch regrowth into the spinal cord. Glia. 2024 Oct;72(10):1766–1784. doi: 10.1002/glia.24581. Epub 2024 Jun 22. PMID: 39141572; PMCID: PMC11325082.

54. Yu, J.A., Castranova, D., Pham, V.N., Weinstein, B.M. (2015) Single cell analysis of endothelial morphogenesis in vivo. Development (Cambridge, England). 142(17):2951–61.

55. Zolessi FR, Poggi L, Wilkinson CJ, Chien CB, Harris WA. Polarization and orientation of retinal ganglion cells in vivo. Neural Dev. 2006 Oct 13;1:2. doi: 10.1186/1749-8104-1-2. PMID: 17147778; PMCID: PMC1636330.

56. Oehlers SH, Cronan MR, Scott NR, Thomas MI, Okuda KS, Walton EM, Beerman RW, Crosier PS, Tobin DM. Interception of host angiogenic signalling limits mycobacterial growth. Nature. 2015 Jan 29;517(7536):612-5. doi: 10.1038/nature13967. Epub 2014 Nov 24. PMID: 25470057; PMCID: PMC4312197.

57. Ellett F, Pase L, Hayman JW, Andrianopoulos A, Lieschke GJ. mpeg1 promoter transgenes direct macrophage-lineage expression in zebrafish. Blood. 2011 Jan 27;117(4):e49–56. doi: 10.1182/blood-2010-10-314120. Epub 2010 Nov 17. PMID: 21084707; PMCID: PMC3056479.

58. Sharrock AV, Mulligan TS, Hall KR, Williams EM, White DT, Zhang L, Emmerich K, Matthews F, Nimmagadda S, Washington S, Le KD, Meir-Levi D, Cox OL, Saxena MT, Calof AL, Lopez-Burks ME, Lander AD, Ding D, Ji H, Ackerley DF, Mumm JS. NTR 2.0: a rationally engineered prodrug-converting enzyme with substantially enhanced efficacy for targeted cell ablation. Nat Methods. 2022 Feb;19(2):205–215. doi: 10.1038/s41592-021-01364-4. Epub 2022 Feb 7. PMID: 35132245; PMCID: PMC8851868.

