## Supplementary figures and images for "Radial astroglia cooperate with microglia to clear neuronal cell bodies during zebrafish optic tectum development"

### Figure S1

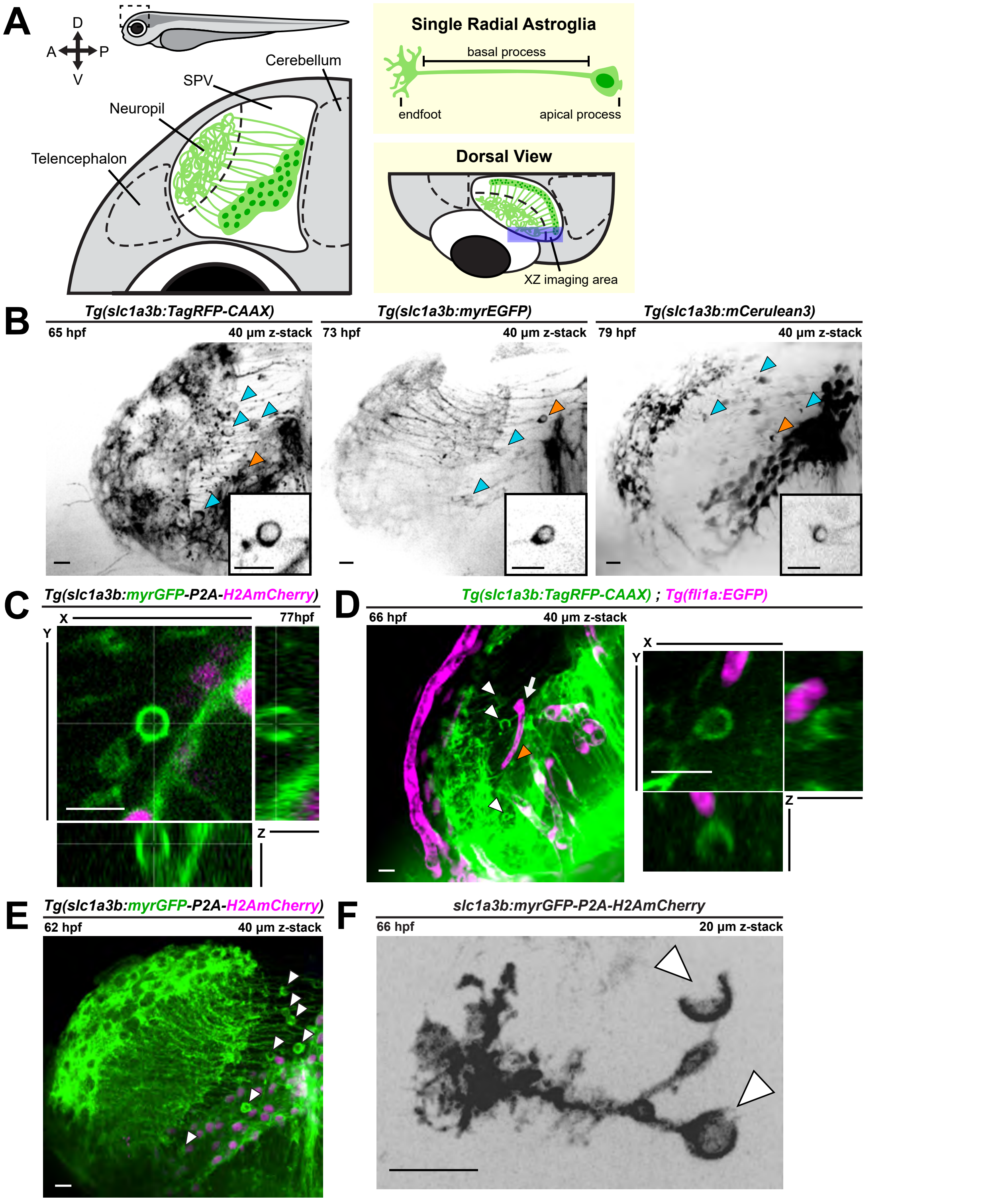

### Figure S2

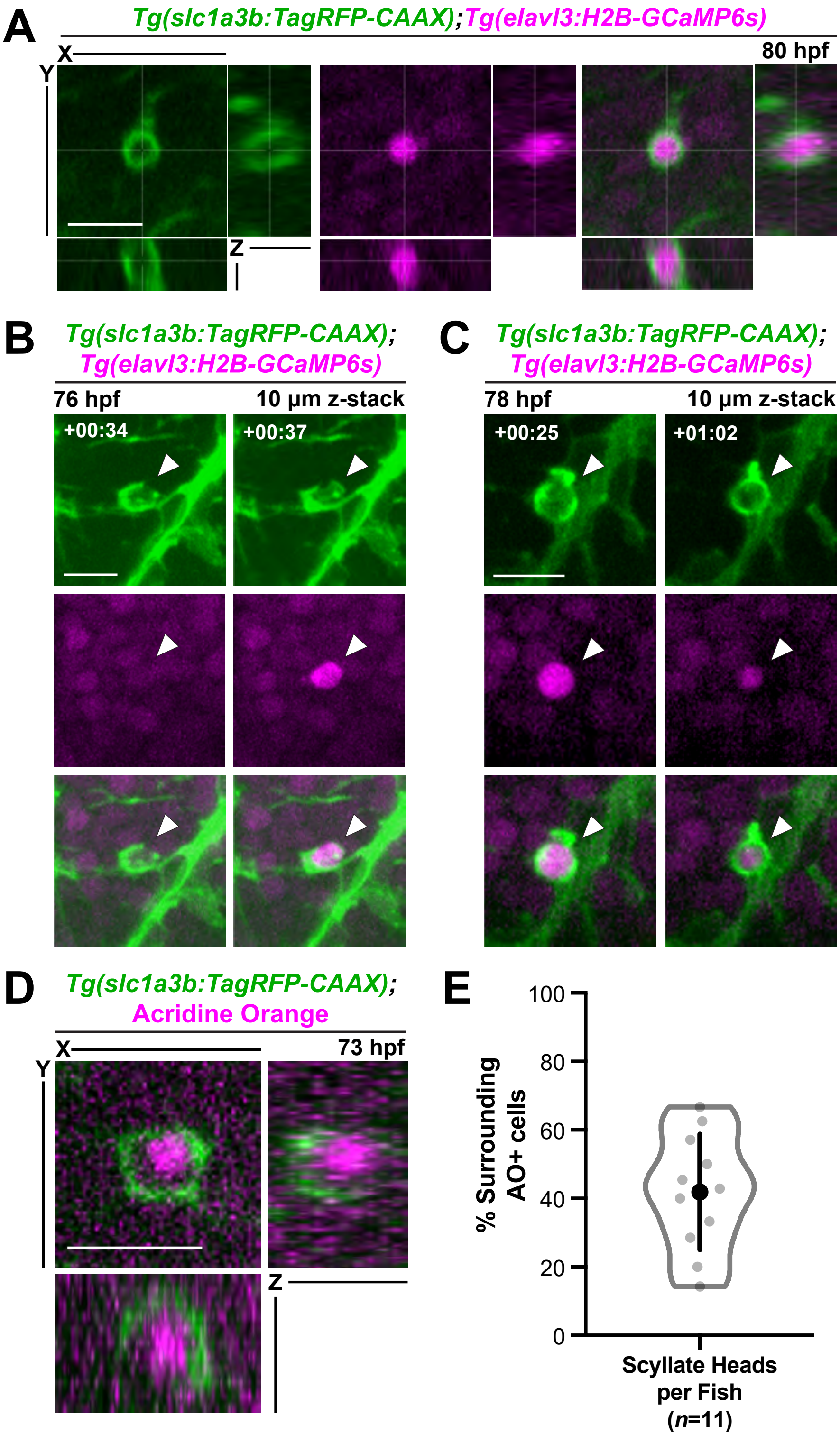

### Figure S3

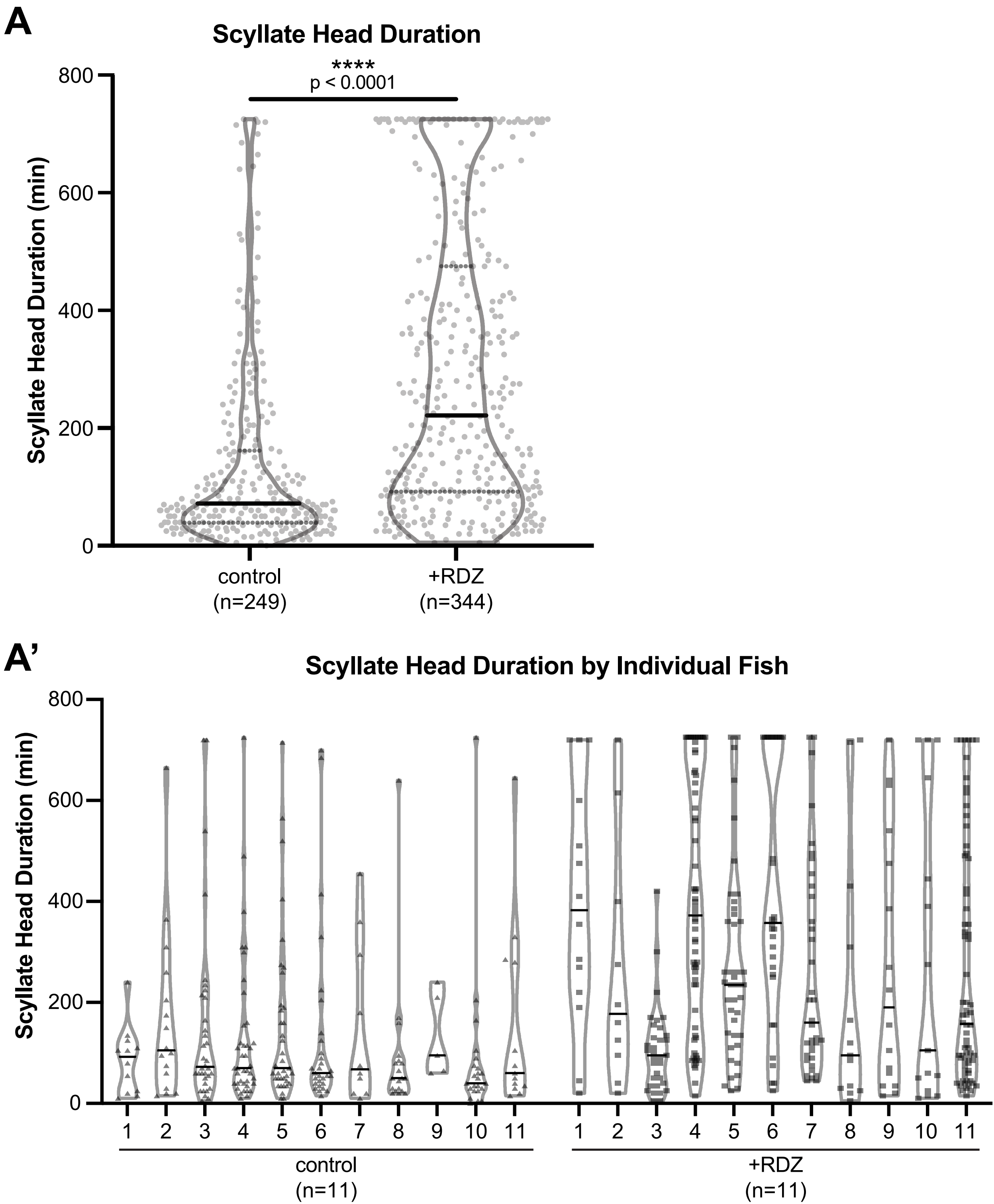
